# Patterns of subregional cerebellar atrophy across epilepsy syndromes: An ENIGMA-Epilepsy study

**DOI:** 10.1101/2023.10.21.562994

**Authors:** Rebecca Kerestes, Andrew Perry, Lucy Vivash, Terence J. O’Brien, Marina K.M. Alvim, Donatello Arienzo, Ítalo K. Aventurato, Alice Ballerini, Gabriel F. Baltazar, Núria Bargalló, Benjamin Bender, Ricardo Brioschi, Eva Bürkle, Maria Eugenia Caligiuri, Fernando Cendes, Jane de Tisi, John S. Duncan, Jerome P. Engel, Sonya Foley, Francesco Fortunato, Antonio Gambardella, Thea Giacomini, Renzo Guerrini, Gerard Hall, Khalid Hamandi, Victoria Ives-Deliperi, Rafael B. João, Simon S. Keller, Benedict Kleiser, Angelo Labate, Matteo Lenge, Cassandra Marotta, Pascal Martin, Mario Mascalchi, Stefano Meletti, Conor Owens-Walton, Costanza B. Parodi, Saül Pascual-Diaz, David Powell, Jun Rao, Michael Rebsamen, Johannes Reiter, Antonella Riva, Theodor Rüber, Christian Rummel, Freda Scheffler, Mariasavina Severino, Lucas S. Silva, Richard J. Staba, Dan J. Stein, Pasquale Striano, Peter N. Taylor, Sophia I. Thomopoulos, Paul M. Thompson, Domenico Tortora, Anna Elisabetta Vaudano, Bernd Weber, Roland Wiest, Gavin P. Winston, Clarissa L. Yasuda, Hong Zheng, Carrie R. McDonald, Sanjay M. Sisodiya, Ian H. Harding, the ENIGMA-Epilepsy Working Group

## Abstract

**Objective:** The intricate neuroanatomical structure of the cerebellum is of longstanding interest in epilepsy, but has been poorly characterized within the current cortico-centric models of this disease. We quantified cross-sectional regional cerebellar lobule volumes using structural MRI in 1,602 adults with epilepsy and 1,022 healthy controls across twenty-two sites from the global ENIGMA-Epilepsy working group.

**Methods:** A state-of-the-art deep learning-based approach was employed that parcellates the cerebellum into 28 neuroanatomical subregions. Linear mixed models compared total and regional cerebellar volume in i) all epilepsies; ii) temporal lobe epilepsy with hippocampal sclerosis (TLE-HS); iii) non-lesional temporal lobe epilepsy (TLE-NL); iv) genetic generalised epilepsy; and (v) extra-temporal focal epilepsy (ETLE). Relationships were examined for cerebellar volume versus age at seizure onset, duration of epilepsy, phenytoin treatment, and cerebral cortical thickness.

**Results:** Across all epilepsies, reduced total cerebellar volume was observed (*d*=0.42). Maximum volume loss was observed in the corpus medullare (*d*_max_=0.49) and posterior lobe grey matter regions, including bilateral lobules VIIB (*d*_max_= 0.47), Crus I/II (*d*_max_= 0.39), VIIIA (*d*_max_=0.45) and VIIIB (*d*_max_=0.40). Earlier age at seizure onset (*ηρ*^2^_max_=0.05) and longer epilepsy duration (*ηρ*^2^_max_=0.06) correlated with reduced volume in these regions. Findings were most pronounced in TLE-HS and ETLE with distinct neuroanatomical profiles observed in the posterior lobe. Phenytoin treatment was associated with reduced posterior lobe volume. Cerebellum volume correlated with cerebral cortical thinning more strongly in the epilepsy cohort than in controls.

**Significance:** We provide robust evidence of deep cerebellar and posterior lobe subregional grey matter volume loss in patients with chronic epilepsy. Volume loss was maximal for posterior subregions implicated in non-motor functions, relative to motor regions of both the anterior and posterior lobe. Associations between cerebral and cerebellar changes, and variability of neuroanatomical profiles across epilepsy syndromes argue for more precise incorporation of cerebellum subregions into neurobiological models of epilepsy.

**Key points:**

- Cerebellar involvement in epilepsy is poorly understood within current cortico-centric models of this disease
- We used a novel, deep learning segmentation tool to parcellate the cerebellum into 28 anatomical subunits using an international MRI dataset of 1,602 individuals with epilepsy (aged between 18 and 65 years old) including temporal lobe epilepsy with hippocampal sclerosis (TLE-HS, *n*=562), TLE non-lesional (TLE-NL, *n*=284), generalised genetic epilepsy (GGE, *n*=186) and extra temporal focal epilepsy (ETLE, *n*=251) and 1,022 controls.
- Across all epilepsies (vs. controls) robust changes in the corpus medullare and posterior lobe “non-motor” regions were observed, with maximal differences in bilateral VIIB and Crus II lobules. Lower volume of these regions correlated with longer disease duration. Anterior “motor lobe” regions were relatively spared.
- Findings were most pronounced in TLE-HS and ETLE groups, with distinct neuroanatomical profiles observed.
- Cortical thinning was associated with pronounced cerebellar volume loss in TLE-HS epilepsy, relative to controls.

**ETHICAL PUBLICATION STATEMENT:** We confirm that we have read the Journal’s position on issues involved in ethical publication and affirm that this report is consistent with those guidelines.

## INTRODUCTION

Epilepsy is a prevalent, chronic group of neurological diseases that affects over 70 million people worldwide^1^, and encompasses many different disorders, of which temporal lobe epilepsy (TLE) is the most common in adults^2^. Current models of the disease conceptualise epilepsy as involving widespread cortical and subcortical network disturbances, including reduced cortical thickness and white matter abnormalities.^3,4^ As such, studies have frequently overlooked the cerebellum. However, a plethora of evidence from electrophysiological and optogenetic studies in animals,^5–7^ as well as non-invasive imaging studies in humans^8,9^ provide evidence for a role of the cerebellum in seizure generation. Further, atrophy of the cerebellum is associated with poorer prognosis following therapeutic temporal resection.^10^ These findings highlight the cerebellum as a potential target for therapeutic intervention in epilepsy and underscore the importance of incorporating the cerebellum into neurobiological models of epilepsy.

Structural magnetic resonance imaging (MRI) has increasingly been used to localise and quantify cerebellar grey and white matter volume changes in people with epilepsy. The most consistent finding has been reduced total cerebellar grey matter volume (GMV) in people with TLE, compared to controls.^9,11–16^ Many studies suggest a negative correlation between total cerebellar volume and duration of epilepsy^11–15^ and earlier age at seizure onset.^11,17^ Bilateral reduction of cerebellar GMV in TLE regardless of the side of seizure focus has been reported by some authors,^14,18–20^ while others have found GMV cerebellar changes most pronounced ipsilateral to the side of the epileptogenic focus.^21,22^ Evidence for changes in cerebellar white matter has been inconsistent.^15,16,23,24^

Collectively these studies have highlighted the presence of global cerebellar atrophy in epilepsy. However, these studies are limited as they have largely neglected the cerebellum’s distinct underlying anatomical subregions (lobules) and associated functional heterogeneity. In addition, these studies have included modest sample sizes and typically have been confined to one epilepsy syndrome, limiting a thorough investigation of cerebellar subregions across epilepsy syndromes. Anatomically, the cerebellum is divided along its superior to inferior axis into three lobes: anterior, posterior (further divided into superior and inferior divisions) and flocculonodular.^25^ Only a few studies have distinguished the cerebellar anterior and posterior lobes in epilepsy,^10,15,16^ and largely report GMV reductions localised to the posterior lobes. Only one study has quantified cerebellar volume of 17 subregions and reported evidence for increased vermal and decreased posterior lobule volumes, in TLE.^10^ This work provides an initial indication that cerebellar changes in people with epilepsy are likely spatially non-uniform, supporting the need for larger, more spatially precise examinations of cerebellar atrophy across different epilepsy syndromes.

The development of new machine learning based approaches for optimised and automated feature-based parcellation of the cerebellum allows for more spatially precise mapping of cerebellar anatomy. These approaches are superior to voxel-based approaches (including voxel-based morphometry; VBM) for accurately quantifying cerebellar volume at the lobular level.^26^ One such approach, called Automatic Cerebellum Anatomical Parcellation using U-Net with Locally Constrained Optimization (ACAPULCO), uses a deep learning algorithm to automatically parcellate the cerebellum into 28 anatomical subunits.^27^ In contrast to registration-based approaches such as VBM, feature-based segmentation ACAPULCO performs on par with leading approaches for automatic cerebellar parcellation, has broad applicability to both healthy and atrophied cerebellums, and is more time-efficient than other approaches.^27^

Here, in the largest study of cerebellar volume in epilepsy to date, we applied ACAPULCO to quantify cerebellar lobule volumes from 1602 individuals with epilepsy and 1022 healthy controls from the global ENIGMA-Epilepsy working group. We undertook multisite mega-analyses to infer regional cerebellar volumetric differences in: (i) all epilepsies; (ii) hippocampal sclerosis-related TLE, considering left (TLE-HS-L) and right-sided (TLE-HS-R) sclerosis as independent groups; (iii) non-lesional TLE (TLE-L-NL and TLE-R-NL); (iv) genetic generalised epilepsy (GGE), and (v) extra-temporal focal epilepsy (ETLE), compared to controls. As a secondary aim, epilepsy syndromes were directly compared, to assess for syndrome-specific changes in total and regional cerebellar volume. Relationships between cerebellar regional volume, age at seizure onset, and epilepsy duration were examined. Finally, given reports of a relationship between chronic phenytoin treatment and cerebellar atrophy^12,28^, we examined associations between cerebellar volume and phenytoin treatment.

## MATERIALS AND METHODS

### Study sample

Sixteen sites were included in this cross-sectional study, totalling 1,602 adult people with epilepsy and 1,022 age-and sex-matched controls. Demographic and clinical characteristics of the sample are listed in **Tables 1** and **S1**. Exclusion criteria included participants with a comorbid progressive disease (e.g., Rasmussen’s encephalitis) or MRI-visible lesions in the cerebellum; however, patients with supratentorial cortical dysplasias-related epilepsy or cortical lesions, were not excluded. An epilepsy specialist assessed seizure and syndrome classifications at each centre, using International League Against Epilepsy terminology^29,30^. For TLE, all individuals presenting with the typical electroclinical constellation and clinical semiology of this disorder were included.^30^ For the TLE-HS subgroups, people with unilateral HS had a neuroradiologically confirmed diagnosis of hippocampal atrophy and increased T2 signal on an epilepsy protocol clinical MRI. TLE-L and TLE-R disorders without HS (as indicated by a normal MRI) were considered as independent groups (TLE-L-NL and TLE-R-NL, respectively). For GGE, people with juvenile onset myoclonic epilepsy, absence or myoclonic seizures with generalized spike-wave discharges on EEG, were included in this group. The ETLE group included people with epileptogenic foci identified outside of the temporal lobe. All remaining epilepsies that could not be classified into the pre-defined syndrome groups were combined into an “all other epilepsies” group (**Table S2**).

**Table 1.** Demographic and clinical characteristics of the total sample.

|  | *N* | Age (SD) | Sex, % female | Age of onset (SD) | Duration epilepsy (SD) |
| --- | --- | --- | --- | --- | --- |
| Controls | 1022 |  |  |  |  |
| All epilepsies | 1602 | 36.2 (11.8) | 56.5 | 18.3 (13.6) | 17.5 (14.1) |
| TLE-HS-L | 320 | 38.8 (10.8) | 54 | 13.8 (11.5) | 25.1 (14.3) |
| TLE-HS-R | 242 | 40.3 (10.9) | 57 | 15.2 (12.1) | 24.6 (14.7) |
| TLE-NL-L | 153 | 37.8 (12.5) | 63 | 26.0 (15.3) | 12.2 (11.9) |
| TLE-NL-R | 131 | 37.3 (11.5) | 59 | 26.5 (13.7) | 10.3 (10.3) |
| GGE | 186 | 29.2 (9.7) | 66 | 13.4 (8.3) | 15.4 (10.5) |
| ETLE | 251 | 32.1 (10.8) | 54 | 15.1 (10.6) | 16.5 (12.1) |
| All other epilepsies | 317 | 36.4 (12.3) | 50 | 24.2 (15.1) | 12.5 (11.0) |

**Table 1. Demographic and clinical characteristics of the total sample**
|  | *n* | Age (SD) | Sex, % female | Age of onset (SD) | Duration epilepsy (SD) |
| --- | --- | --- | --- | --- | --- |
| Controls | 1022 |  |  |  |  |
| All epilepsies | 1602 | 36.2 (11.8) | 56.5 | 18.3 (13.6) | 17.5 (14.1) |
| TLE-HS-L | 320 | 38.8 (10.8) | 54 | 13.8 (11.5) | 25.1 (14.3) |
| TLE-HS-R | 242 | 40.3 (10.9) | 57 | 15.2 (12.1) | 24.6 (14.7) |
| TLE-NL-L | 153 | 37.8 (12.5) | 63 | 26.0 (15.3) | 12.2 (11.9) |
| TLE-NL-R | 131 | 37.3 (11.5) | 59 | 26.5 (13.7) | 10.3 (10.3) |
| GGE | 186 | 29.2 (9.7) | 66 | 13.4 (8.3) | 15.4 (10.5) |
| ETLE | 251 | 32.1 (10.8) | 54 | 15.1 (10.6) | 16.5 (12.1) |
| All other epilepsies | 317 | 36.4 (12.3) | 50 | 24.2 (15.1) | 12.5 (11.0) |

### Image processing and analysis: ACAPULCO

Whole-brain, T1-weighted MR images were collected from each participant. Scanner descriptions and acquisition protocols for all sites are reported in **Table S3**. We treated each individual scanner and/or data acquisition protocol used in the collection of MRI scans as a separate ‘site’ during statistical analysis (see below). Each image was processed in accordance with the ENIGMA cerebellum parcellation protocol, as fully described elsewhere^31^ (https://enigma.ini.usc.edu/protocols/imaging-protocols/). In brief, the cerebellum was parcelled into 28 subregions (left and right lobules I-III, IV, V, VI, Crus I, Crus II, VIIB, VIIIA, VIIIB, IX and X; bilateral vermis VI, VII, VIII, IX and X, and bilateral corpus medullare (central white matter and deep cerebellar nuclei) using the deep learning-based algorithm ACAPULCO^27^ (version 0.2.1; https://gitlab.com/shuohan/acapulco). Outputs were quality checked by visual inspection followed by quantitative identification of outlier volumes that were greater or less than 2.698 standard deviations from the group mean.

### Statistical analysis

All statistical analyses of cerebellar volume were carried out using R version 4.1.0.^32^ We fit linear mixed effects regression models (LMM) using *lme4* and *lmerTest* packages in R, with diagnosis (Dx; i.e., control or epilepsy), age, age^2^, sex and intracranial volume (ICV) as fixed factors and site as a random factor. Age^2^ was included given evidence of non-linear effects of age on brain volume loss in the cerebellum and cerebral cortex.^33^ Model (1) was fit for total cerebellar volume (sum of all 28 cerebellar regions of interest (ROI)) and each cerebellar lobule individually and repeated for the total sample (all epilepsy vs controls) and each subgroup of interest: TLE-HS-L, TLE-HS-R, TLE-L-NL, TLE-R-NL, GGE and ETLE.

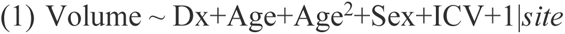

LMMs were repeated to compare epilepsy syndromes to one another. For all analyses, results were FDR (*P*<0.05) corrected for multiple comparisons. Cohen’s *d* effect sizes with 95% confidence intervals were calculated for each of the ROIs, based on the estimated marginal means and Satterthwaite’s approximation for degrees of freedom.^34^ Positive effect size values correspond to people with epilepsy having *lower* values relative to controls.

### Regression with clinical variables and phenytoin therapy

LMMs were used to test for associations between each ROI volume (and total cerebellar volume) and i) duration of epilepsy and ii) age at onset in all epilepsies, as well as in each epilepsy disorder subtype. Partial η^2^ (eta-squared) is reported as a measure of effect size. To investigate the association between phenytoin treatment and volume of each ROI in all epilepsies, a binary variable indicating exposure to phenytoin (“Yes” = have been treated with phenytoin at some point for any duration or “No” = never been treated with phenytoin) was treated as the predictor of interest, with age, age^2^, duration of epilepsy and ICV included as fixed factors and site as a random factor. To examine site-specific effects, we also fit LMMs separately for each site.

### Laterality

In addition, we investigated whether the side of seizure focus in patients with TLE (e.g., TLE-HS-L vs TLE-HS-R) was associated with left/right asymmetry of cerebellar volumes. Methodological details can be found in the supplementary material.

## RESULTS

### Participant demographics

A one-way ANOVA with group as a fixed effect revealed differences across the eight groups in age [*F*(7,2614)=27.76, *P*<0.001]. Two one-way ANOVAs across the seven patient groups revealed group differences in age at seizure onset [*F*(6,1235)=37.12, *P*<0.001] and duration of epilepsy [*F*(6,1248)=45.06, *P*<0.001]. Post *hoc* comparisons revealed that controls were younger than the TLE-L, TLE-R, TLE-L-NL, and ‘all other’ epilepsy groups, but significantly older than the GGE group (*P*<0.05). The TLE-L-NL and TLE-R-NL groups and ‘all other’ epilepsy groups had a later age of seizure onset compared to all other syndromes (*P*<0.05). In addition, TLE-HS-L and TLE-HS-R had a longer duration of epilepsy compared to all other groups.

Four of the sixteen data collection sites used more than one imaging sequence across their cohort; these sites were also broken into sub-sites, resulting in a total of 22 sites included in our mixed linear regression models.

### All epilepsies group

#### All epilepsies versus healthy controls

Compared to controls, people with epilepsy had significantly reduced total cerebellar (grey and corpus medullare) volume (*d=*0.42, 95% [0.33, 0.52]). ROI analyses revealed significantly reduced volume of the corpus medullare (*d*=0.49, 95% [0.39, 0.58]), and reduced grey matter volume in 21 cerebellar lobules, with small to moderate effect sizes (*d*_min_=0.11, *d*_max_=0.40, all *P*<0.05 FDR; **Fig 1**). The largest effect sizes were observed in predominantly bilateral superior and inferior posterior lobe regions including left and right VIIB, right and left Crus II, right V, left VIIIB, left Crus I and right VIIIB. There were no significant between-group differences for the remaining cerebellar lobules (**Fig 1, Table S4**).

**Fig 1.**
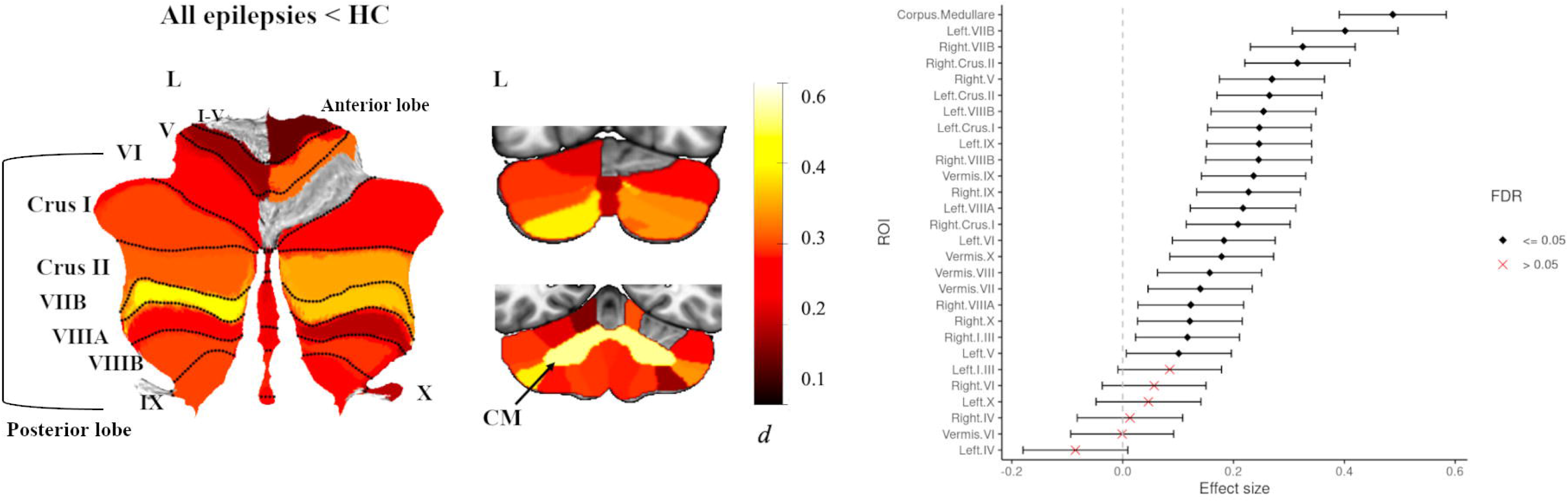
Atlas-based effect size (Cohen’s *d*) maps, MNI-based coronal slices (top: y= -72; bottom: y= -54) and forest plots (Cohen’s *d* +/- 95% confidence interval) of the significant between-group differences for all epilepsies vs. healthy controls (HC). Note: positive effect sizes reflect epilepsy patients < HC. Regions significant at *P* < 0.05 FDR corrected are depicted in colour (see Table S4 for full tabulation).

#### Duration of epilepsy and age at seizure onset

Longer duration of epilepsy corresponded with smaller total cerebellar volume across all epilepsies (partial η^2^=0.032, 95% [0.017, 0.061], *P*<0.001); **Fig 2a**. Analyses of individual cerebellar subregions revealed significant negative correlations between duration of epilepsy and regions that also showed reduced grey matter volume (all *P*<0.05 FDR; **Table S9**). Earlier age at onset corresponded with smaller total cerebellar volume (partial η^2^=0.024, 95% [0.051, 0.011], *P*<0.001); **Fig 2b**. Nineteen cerebellar subregions showed a significant positive association with age at disease onset. The strongest effects were seen for regions that showed reduced grey matter in people with epilepsy (vs. controls) (all *P*<0.05 FDR; **Table S10**).

**Fig 2.**
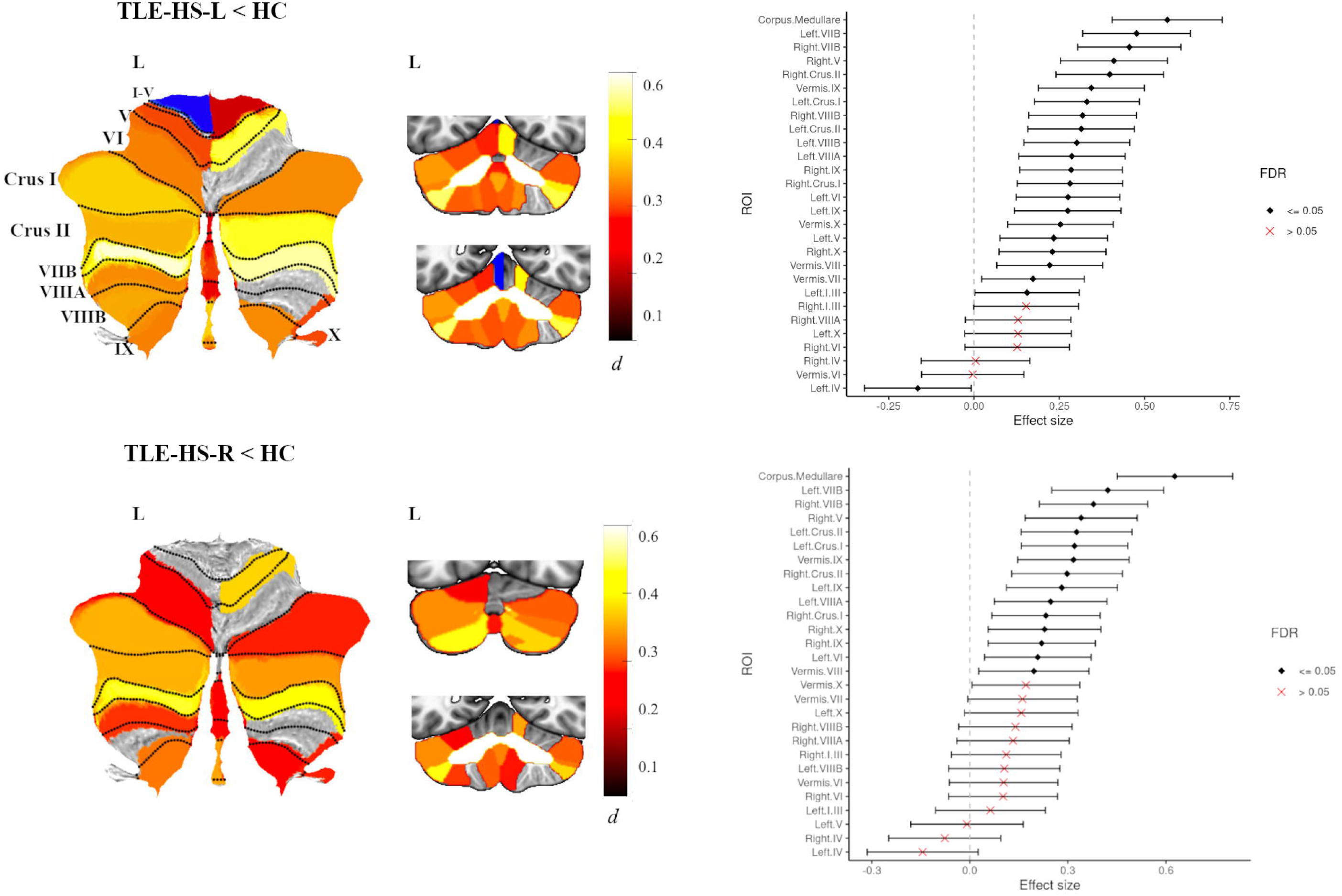
Scatter-plots showing the association between a) duration of illness and b) age at seizure onset, and total cerebellar volume in epilepsy patients, *P* < 0.001.

### TLE-HS group

#### TLE-HS versus healthy controls

In TLE-HS-L (*n*=320) there was significantly reduced total cerebellar volume (*d=*0.56, 95% [0.40, 0.71]). ROI analyses revealed significantly reduced volume of the corpus medullare compared to healthy controls (*d=*0.56, 95% [0.40, 0.72]). In addition, the TLE-HS-L group also showed reduced grey matter volume in 22 cerebellar lobules, with small to moderate effect sizes (*d*_min_=0.15, *d*_max_=0.48, all *P*<0.05 FDR; **Fig 3**), with the largest effect sizes seen in left and right VIIB, right Crus II and right V. The TLE-HS-L group also showed increased cerebellar volume in left IV (*d=-*0.16, 95% [-0.32, -0.01]). There were no significant between-group differences for the remaining cerebellar lobules (**Fig 3, Table S5**).

**Fig 3.**
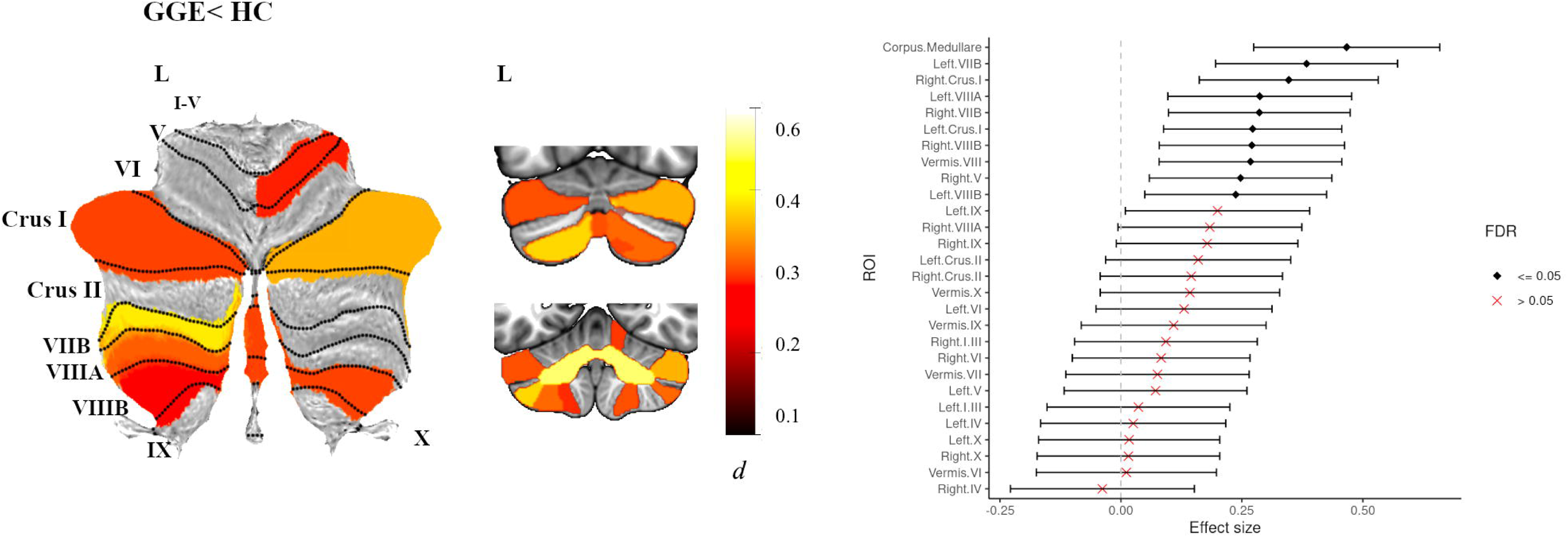
Atlas-based effect size (Cohen’s *d*) maps, MNI-based coronal slices (top: y= -72; bottom: y= -54) and forest plots (Cohen’s *d* +/- 95% confidence interval) of the significant between-group differences for TLE-HS-L (top panel) and TLE-HS-R (lower panel) vs. healthy controls (HC). Positive effect sizes reflect TLE-HS-L and TLE-HS-R < HC respectively. Regions significant at *P* < 0.05 FDR corrected are depicted in colour (red-yellow for epilepsy < HC; blue for epilepsy > HC); see Table S5 for full tabulation).

In TLE-HS-R (*n*=242) total cerebellar volume was significantly reduced (*d=*0.48, 95% [0.30, 0.64]). ROI analyses significantly reduced volume of the corpus medullare compared to healthy controls (*d=*0.63, 95% [0.45, 0.80]) (**Fig 3**). However, the TLE-HS-R group showed less pronounced (compared to TLE-HS-L group) cerebellar volume loss across individual cerebellar lobules. Reduced grey matter was found for 14 cerebellar lobules (*d*_min_=0.19, *d*_max_=0.42, all *P*<0.05 FDR; **Fig 3**), with the largest effect sizes seen for left and right VIIB, left Crus I and right V (**Table S5**).

#### Duration of epilepsy and age at seizure onset

There were no significant relationships between duration of epilepsy or age at onset and total cerebellar volume in the TLE-HS-L group (**Tables S9, S10**). In the right TLE-HS group however, longer duration of epilepsy corresponded with smaller total cerebellar volume (partial η^2^=0.06, 95% [0.01, 0.15], *P=*0.010; **Table S9**); in addition, later age at disease onset corresponded with higher total cerebellar volume (partial η^2^=0.06, 95% [0.01, 0.14], *P*=0.016; **Table S10**).

### TLE non-lesional group

#### TLE non-lesional (NL) versus healthy controls

Compared to healthy controls, the left TLE-NL group (*n*=153) did not show significant differences in total cerebellar volume or individual cerebellar lobule volumes (all *P*>0.05, FDR). The left TLE-NL group did however, show significantly reduced white matter volume of the corpus medullare (*d=* 0.58, 95% [0.37, 0.79]) (**Fig S1a, Table S6**).

The right TLE-NL group (*n*=131) did not show significant differences in total cerebellar volume, compared to healthy controls (*P*>0.05, FDR), however they showed reduced volume of the corpus medullare (*d=*0.57, 95% [0.33, 0.80]). Right TLE-NL also showed significantly increased grey matter volume of left IV (*d=-*0.38, 95% [-0.61, -0.15]; **Fig S1b, Table S6**.

#### Duration of epilepsy and age at seizure onset

There were no significant relationships between duration of epilepsy or age at disease onset and total cerebellar volume in the TLE NL groups (all *P*>0.05; **Tables S9, S10**).

### GGE group

People with GGE (*n=*186) showed significantly reduced total cerebellar volume (*d=*0.42, 95% [0.23, 0.61]) and reduced volume of the corpus medullare (*d=*0.47, 95% [0.27, 0.66]) compared to healthy controls. Additionally, they showed reduced grey matter volume in 9 cerebellar lobules with small to moderate effect sizes (*d*_min_=0.24, *d*_max_=0.38, all *P*<0.05 FDR; **Fig 4**). Differences were seen for predominantly bilateral posterior regions including left and right VIIB, right Crus I and left VIIIA and right VIIIB (**Fig 4, Table S7**).

**Fig 4.**
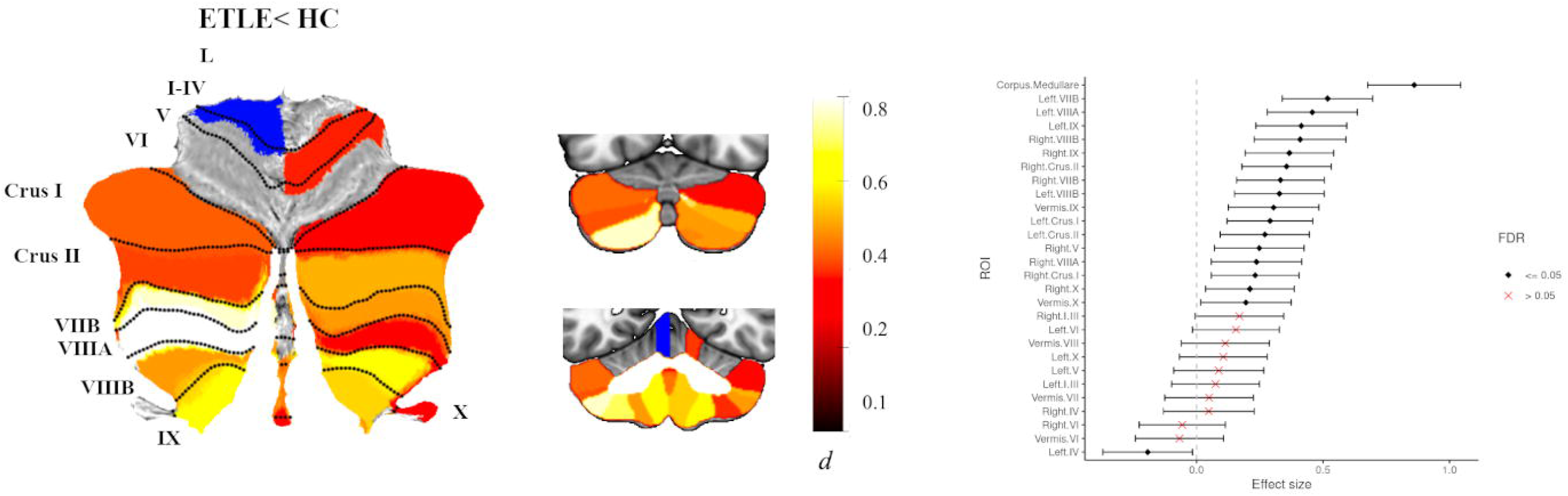
Atlas-based effect size (Cohen’s *d*) maps, MNI-based coronal slices (top: y= -72; bottom: y= -54) and forest plots (Cohen’s *d* +/- 95% confidence interval) of the significant between-group differences for GGE vs. healthy controls (HC). Note: positive effect sizes reflect epilepsy < HC. Regions significant at *P* < 0.05 FDR corrected are depicted in colour. See Table S7 for full tabulation.

#### Duration of epilepsy and age at seizure onset

There was no significant relationship between duration of epilepsy or age at seizure onset and total cerebellar volume in people with GGE (all *P>*0.05; **Tables S9** and **S10**).

### ETLE group

Individuals with ETLE (*n*=251) showed significantly reduced total cerebellar volume (*d*=0.58, 95% [0.40, 0.76]) and reduced volume of the corpus medullare (*d=*0.86, 95% [0.67, 1.0]), compared to healthy controls. ROI analyses showed reduced grey matter volume in 17 cerebellar lobules with small to moderate effect sizes (*d*_min_=0.12, *d*_max_=0.52, all *P*<0.05 FDR; **Fig 5**). The largest effect sizes were seen for predominantly left-sided posterior non-motor and motor regions: left VIIB, left VIIIA, and left IX. People with ETLE also showed increased grey matter volume of left anterior lobule IV compared to controls (*d=-*0.19, 95% [-0.37, -0.01]) (**Fig 5, Table S8**).

**Fig 5.**
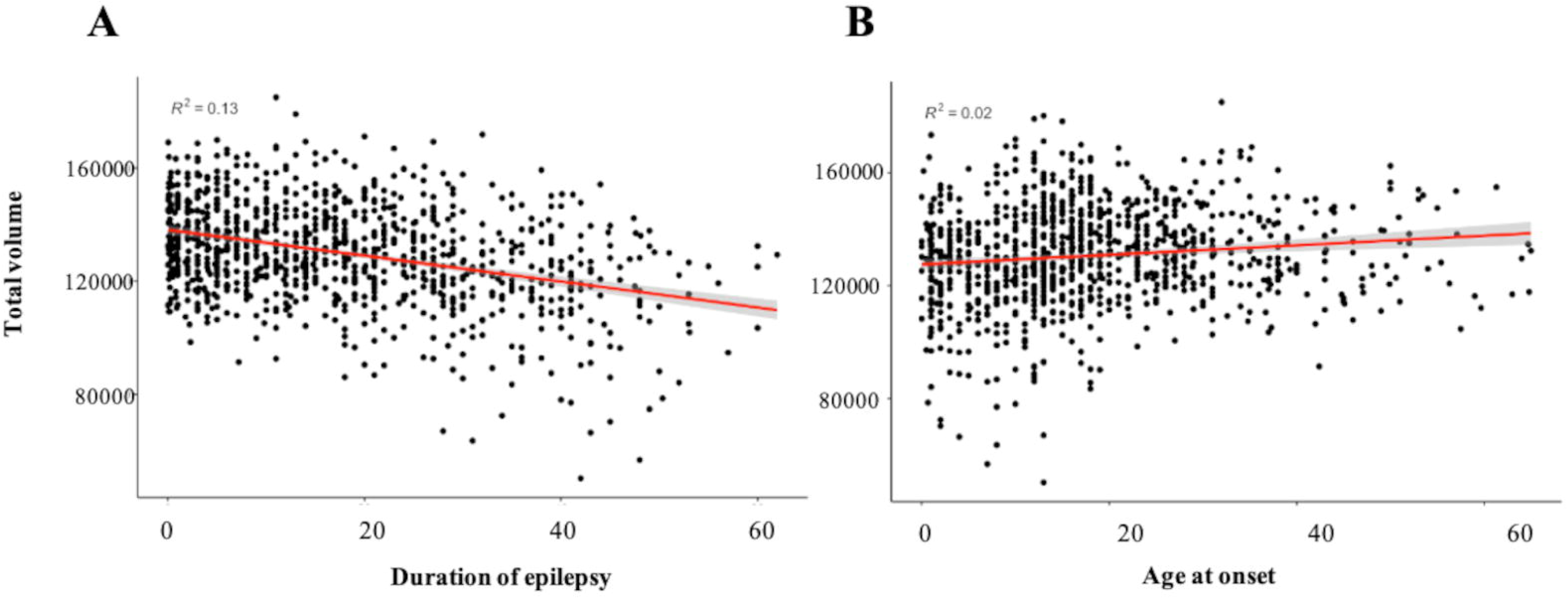
Atlas-based effect size (Cohen’s *d*) maps, MNI-based coronal slices (top: y= -72; bottom: y= -54) and forest plots (Cohen’s *d* +/- 95% confidence interval) of the significant between-group differences for ETLE vs. healthy controls (HC). Note: positive effect sizes reflect epilepsy patients < HC. Regions significant at *P* < 0.05 FDR corrected are depicted in colour (red-yellow for ETLE < HC; blue for ETLE > HC); see Table S8 for full tabulation.

#### Duration of epilepsy and age at seizure onset

Longer duration of epilepsy was associated with smaller total cerebellar volume (partial η^2^=0.05, 95% [0.01, 0.13], *P*<0.05 FDR; **Table S9**). In addition, later age of onset with associated with larger total cerebellar volume (partial η^2^=0.05, 95% [0.01, 0.13], *P*<0.05 FDR; **Table S10**).

### Comparisons Between Epilepsy Syndromes

The most significant differences were observed between the TLE-HS-L and TLE-NL-L groups in posterior lobe regions, where the TLE-HS-L group had significantly lower right VIIB (d=0.51, 95% [0.20, 0.77]) and right Crus II (d=0.39, 95% [0.11, 0.67]) volumes. The ETLE group showed significantly lower volume of right VIIB (d=0.60, 95% [0.20, 0.89]), left and right lobule IX (d=0.40, 95% [0.14, 0.65] and d=0.46, 95% [0.16, 0.76] respectively), compared to the TLE-NL-L group. Finally, compared to the GGE group, people with ETLE showed significantly decreased volume of left IX (d=0.28, 95% [0.10, 0.50]).

### Relationships between Cerebellar and Cerebral Structural Changes

We explored cerebro-cerebellar relationships to assess whether the magnitude of cerebellar volume loss observed in people with epilepsy, mirrored that seen in the cerebral cortex. Freesurfer-derived cortical thickness measures were available for 1,586 individuals across 8 sites (controls= 804, epilepsies= 782). Methodological details are provided in the supplementary material. Results showed a significant interaction between cortical thickness and diagnosis on predicting cerebellar total volume (p=0.01). Marginal effects plots (controlling for all other covariates), showed that a reduction in mean cortical thickness was associated with a more pronounced decrease in cerebellar volume for those with epilepsy (**Fig S2**). Additionally, we explored the interaction of subdiagnosis with cortical thickness. Of the epilepsy syndromes, a significant interaction between cortical thickness and subdiagnosis was only observed for the right and left TLE-HS cohorts (p=0.003), suggesting that the reduction in cerebellum volume relative to cortical thickness is more pronounced in TLE-HS.

### Volume changes associated with phenytoin therapy

Phenytoin therapy data (*n*=802; Yes=161, No=641) was derived from 7 sites. People treated with phenytoin were all affected by focal epilepsy. In contrast, the phenytoin naïve group consisted of focal, generalised, and unspecified epilepsy cases. We therefore restricted analyses of phenytoin use to focal epilepsies only. Independent sample t-tests showed that people treated with phenytoin had a significantly longer duration of illness than phenytoin-naïve individuals, at four of the sites (**Table S11**). Linear regression controlling for duration of epilepsy, revealed that phenytoin therapy was associated with reduced total cerebellum volume (*d*=0.39, 95% [0.12, 0.63]) and reduced grey matter of 10 posterior lobe regions (all *P*<0.05 FDR; **Fig S3**). *Post hoc* analyses for each site independently showed cerebellar volume loss in phenytoin treated individuals to be associated with medium to large effect sizes for three sites, however there was substantial variability in effect sizes across sites (*d*_min_=0.25, *d*_max_=0.86).

### Laterality

There were no significant associations between laterality of disease (right vs. left) and total cerebellar volume asymmetry, or individual cerebellar lobule asymmetry for any of the epilepsy syndromes (all *P*>0.05).

## DISCUSSION

In the largest quantitative study of cerebellum morphometry in epilepsy to date, we performed a comprehensive assessment of regional cerebellar atrophy in over 1600 adults with epilepsy. We report significant cerebellar volume reductions, principally weighted to the posterior cerebellar lobe. These differences were observed across epilepsy syndromes, although were most pronounced in TLE-HS and ETLE. Smaller volumes in posterior lobe regions were associated with longer duration of epilepsy. Exploration of cerebro-cerebellar volume ratios demonstrated that cortical thinning was associated with a more pronounced decrease in cerebellar volume in individuals with epilepsy relative to controls. Collectively, these findings suggest involvement of the cerebellum in epilepsy and raise important questions about the potential vulnerability of different cerebellar subregions in the causes, consequences and clinical expression of specific disease features in epilepsy.

### Dissociation in anterior ‘motor’ vs posterior ‘non-motor’ cerebellar lobe involvement across epilepsies

The spatial non-uniformity of the observed anatomical changes provides evidence for a dissociation in anterior (motor) vs. posterior (non-motor) lobe involvement in epilepsy. Indeed, previous studies report evidence for reduced superior^15^ and inferior posterior lobe^16^ volume in epilepsy, alongside spared or increased anterior lobe volume. The posterior lobe, particularly lobules VII (Crus I/II and VIIB) where the observed anatomical changes were maximal, are predominantly ‘non-motor’ regions of the cerebellar cortex, and are preferentially connected to prefrontal and posterior parietal regions of the cerebral cortex.^35,36^ Functional mapping studies ascribe these areas to cognitive, language, and attentional processes.^37,38^ Functionally, these regions are also part of the frontoparietal and salience resting state networks that are selectively vulnerable to neurodegeneration in Alzheimer’s disease and related conditions that show evidence of cerebellar functional changes and associated cognitive decline. In contrast, the cerebellar anterior lobe (‘motor cerebellum’), flocculonodular lobe, and vermis were relatively spared. This is perhaps surprising when drawing parallels to *cerebral cortical* thickness changes of cerebral motor regions that are functionally and anatomically connected to motor regions of the cerebellar cortex, in people with epilepsy.^4^ Specifically, reduced bilateral cortical thickness of the pre and postcentral gyri-motor cortical regions connected to anterior lobe and motor regions of the posterior lobe, namely lobule VIII-have been reported in TLE-HS and GGE epilepsy disorders.^4^ It is important to note, that although volumetric changes were maximal in superior posterior (predominantly non-motor) regions, the anterior and inferior posterior (motor/pre-motor) lobes were still significantly affected. Given strong evidence of the cerebellum being a brain region particularly vulnerable to injury and neurodegeneration^39,40^, and our observations of a negative relationship between posterior lobe volume loss and illness duration, we speculate that the posterior lobe may be particularly vulnerable to epilepsy-related cell loss. Clinically, such pathology of the non-motor posterior lobe could be associated with cognitive and emotional co-morbidities in people with epilepsy, and future imaging studies in epilepsy should further explore this possibility.

### Shared and distinct patterns of cerebellar grey and white matter atrophy across epilepsy syndromes

The most robust finding across all epilepsy disorders was reduced corpus medullare volume, suggesting that atrophy of the corpus medullare represents a global/shared feature of the disease. In contrast, regional grey matter atrophy across the cerebellum showed different patterns across epilepsy syndromes, suggesting unique neuroanatomical profiles of cerebellum volume changes. We do, however, refrain from interpreting all volumetric changes in the corpus medullare as unambiguously reflecting white matter atrophy. The dentate (and other deep cerebellar) nuclei cannot be delineated on structural T1w images, therefore the corpus medullare label includes these structures. Nonetheless our findings broadly align with previous investigations that report robust changes to white matter structure,^3^ and more mild changes in grey matter cortical thickness,^4^ in people with TLE.

In contrast to white matter changes, the specific pattern and magnitude of volume changes in the posterior lobe grey matter (and underlying white matter) varied across the subgroups. Reduced grey matter volume was most pronounced for the TLE-HS groups and least pronounced in TLE-NL groups (with the ETLE and GGE groups in-between). TLE is the most common and well characterised subtype of epilepsy^2^ and is associated with unilateral or bilateral hippocampal sclerosis in up to 60-70% of adult cases.^41^ Previous reports showed that even among people with drug resistant TLE, NL individuals have less severe cortical^42,43^ and white matter^3,44^ abnormalities. The current work indicates that this distinction, with its implications, is recapitulated in the cerebellum. Importantly, the TLE-HS group in our study had a more chronic course of illness (longer duration of epilepsy) with a largely childhood/early adolescent onset, compared to NL individuals (**Table 2**). The TLE-HS group also represented the largest group of individuals to be treated with phenytoin in our study. Analyses comparing the lesional (TLE-HS) and non-lesional (NL) groups controlling for duration of illness, showed that TLE-HS-L have marked volume loss of right VIIB and right Crus II that distinguishes them from individuals without HS. The cross-sectional nature of our study, however, precludes us from drawing causal statements on volumetric changes in people with chronic epilepsy. For example, we cannot rule out pre-existing cerebellar brain injury that may have occurred from the initial epileptogenic insult. Moreover, the observed distinctions in cerebellar morphology likely reflect the influences of illness chronicity, phenytoin exposure, neurodevelopmental factors, and/or pathophysiological distinctions in TLE-HS vs NL.

People with GGE showed a similar but much less pronounced pattern of cerebellar grey matter volume reduction compared to the TLE-HS groups, alongside reduced white matter volume as is observed in all epilepsy syndromes. The ETLE group showed a similar magnitude of cerebellum grey matter loss to the TLE-HS and NL groups, but – in an important qualitative distinction-they showed a stronger involvement of left-lateralized ‘motor’ *inferior* posterior cerebellar regions (VIIIA, VIIIB, IX). Notably, our analyses directly comparing epilepsy syndromes (controlling for the duration of disease) showed that left lobule IX volume in the ETLE group was significantly lower than the non-lesional TLE and GGE groups (but not when compared to the TLE-HS groups). Our results suggest that people with ETLE harbour unique cerebellar neuroanatomical features, which are not common to focal epilepsy in general. In summary, our findings point to shared and unique cerebellar anatomical changes across epilepsy disorders.

### Clinical implications

Our findings have clinical implications in the context of advancing treatment in epilepsy. Our findings of an association between posterior cerebellar volume and duration of illness across all epilepsies, in particular, demonstrates a common and potentially progressive neurodegeneration in patients with epilepsy. This was substantiated by our findings of greater cerebellar volume loss relative to cerebral cortical thinning in epilepsy (compared to controls), suggesting the cerebellum is particularly vulnerable to seizure-related atrophy. As recognized by the ILAE, a major challenge for the field is to develop disease-modifying or anti-epileptogenesis treatments.^45^As these medications reach late-stage clinical trials, objective measures of the presence or progress of disease will be needed to support long term claims of disease-modification or anti-epileptogenesis. Posterior cerebellar volume in this case, may be a useful additional biomarker of disease severity and progression. In addition, the cerebellum has advantages in potentially being a generalised marker of continued disease burden, as it is uninfluenced by laterality of seizure onset or the lesion itself. Given the cerebellum has a role in both cognitive and emotional processing,^38^ reduced cerebellar (particularly posterior) volumes could be associated with the common mood and cognitive comorbidities observed in epilepsy. ^46^ If this hypothesised association was found to be correct, cerebellar volumes could be considered when deciding on anti seizure medications (ASMs), with ASMs with substantial known effects on mood or cognition not prescribed as first line therapies for patients with smaller cerebellar volumes.

### Laterality of cerebellar changes across epilepsy

We did not find any significant evidence for lateralized differences in the degree of cerebellar volume loss between the left versus right cerebellum. Notably, however, we did find that right-sided Crus II and VIIB volume in the left TLE-HS group statistically differed from the TLE-NL group, providing some support for crossed-cerebellar changes. Evidence for lateralisation of cerebellar volume asymmetry in epilepsy disorders has been inconsistent with several studies reporting bilateral cerebellar volume reduction, regardless of laterality of seizure onset.^20^ These findings may suggest that cerebellar involvement is not driven by diaschisis via cerebro-cerebellar connectivity, but represents another element of more widespread intracranial derangement associated with the consequences of seizures and their treatment.

### Associations with phenytoin

An association between phenytoin exposure and reduced cerebellar posterior lobe volume in TLE-HS individuals, was also observed. Our study is the most robust and largest evaluation of phenytoin therapy in epilepsy disorders, to date. The pattern of atrophy-notably almost exclusively localized to the posterior lobe-suggests that whilst phenytoin is associated with pronounced targeted cerebellar atrophy, it cannot explain atrophy occurring across the breadth of the cerebellum in these patients and speaks to independent disease-related cerebellar pathology. Whilst our *post hoc* analyses revealed that the findings were driven by three sites that were associated with medium to large effect sizes, the variability in sample sizes across sites for which information on phenytoin therapy was available, may have rendered the analysis underpowered to detect statistically significant effects for smaller sites. Future studies will be required to conduct a more fine-grained assessment of phenytoin-mediated effects on cerebellum atrophy.

### Limitations and Future Considerations

Our study has several limitations. The inconsistency of data availability across sites precluded particular analyses of disease features (e.g., cognitive impairments, comprehensive antiseizure medication history). The GGE and ETLE groups also represented heterogeneous cohorts, perhaps contributing to greater variability and weaker effects compared to the TLE-HS groups. We did not have available information on seizure frequency and severity which precluded analyses of the relationship between cerebellar volume and cumulative seizure burden. Finally, whilst we report strong volumetric changes of the corpus medullare across all epilepsies, we cannot interpret these findings as reflecting pure white matter. Given that the dentate nucleus and other deep cerebellar nuclei cannot be delineated from structural T1w images, these structures were also captured in our segmentation of the corpus medullare. Future studies using quantitative susceptibility mapping are required to rule-out or rule-in volumetric changes in the dentate nucleus as a feature of epilepsy.

In summary, we provide evidence of shared and unique cerebellar atrophy profiles in a large, international cohort of people with epilepsy across epilepsy syndromes. Shared atrophy was observed in the corpus medullare and bilateral posterior lobes, with the strongest effects in the TLE-HS groups, whereas volume loss in other regions (i.e., VIIIA, VIIIB, IX) were only observe in patients with ETLE. Importantly, cerebellar atrophy was associated with longer disease duration and phenytoin therapy, raising concerns for a neurodegenerative aspect to this loss. Future studies examining cerebro-cerebellar spatial covariance will extend upon this work and provide insights to the prominence of cerebellar changes within current cortico-centric neurobiological models of the disease.

## DATA AVAILABILITY STATEMENT

The data that support the findings of this study are available on request from the corresponding author. The data are not all publicly available in a repository as they contain information that could compromise the privacy of research participants. Although there are data sharing restrictions imposed by (i) ethical review boards of the participating sites, and consent documents; (ii) national and trans-national data sharing laws, such as GDPR; and (iii) institutional processes, some of which require a signed MTA for limited and predefined data use, we welcome sharing data with researchers, requiring only that they submit an analysis plan for a secondary project to the leading team of the Working Group (http://enigma.ini.usc.edu).

## AUTHOR CONTRIBUTIONS

R.K., A.P., C.R.M., S.M.S. and I.H.H. contributed to the conception and design of the study & methods. M.K.M.A., A.B., G.F.B., N.Ba., B.Ben., R.B., J.S.D., J.E.Jr., F.F., A.G., R.G., V.I., R.B.J., S.S.K., B.K., A.L., P.M., M.M., S.M., C.P., J.Ra., M.R., A.R., T.R., C.R., M.S., L.S.S., R.J.S., P.N.T., D.T., A.V., G.P.S and C.L.Y. contributed to the imaging data collection. R.K., D.A., I.K.A., A.B., B.B., E.B., M.C., S.F., F.F., G.H., V.I., R.B.J., S.S.K., M.L., C.O.W., C.P., S.P.D., M.R., J.Re., C.R., F.S., R.J.S., P.N.T., A.V., C.L.Y. and H.Z. contributed to imaging data analysis. F.C., A.G., R.G., K.H., S.S.K., A.L., P.M., S.M., T.R., R.J.S., P.S., B.W., G.P.S, R.W., C.R.M. and S.M.S were principle investigators at each site. R.K., A.P., D.P. and I.H.H. were the core analysts and R.K. and I.H.H were the core writing group. A.P., T.O., L.V., N.Ba., B.B., E.B., J.S.D., F.F., S.S.K., B.K., M.M., M.S., R.J.S., P.S., P.N.T., S.I.T., P.M.T., D.T., G.P.S, C.L.Y., C.R.M. and S.M.S contributed to editing of the manuscript.

## PATIENT CONSENT STATEMENT

All study participants provided written informed consent for the local study, and the local institutional review boards and ethics committees approved each included cohort study.

## REFERENCES

1. Thijs RD, Surges R, O’Brien TJ, Sander JW. Epilepsy in adults. Lancet. Feb 16 2019;393(10172):689–701. doi:10.1016/s0140-6736(18)32596-0

2. Téllez-Zenteno JF, Hernández-Ronquillo L. A review of the epidemiology of temporal lobe epilepsy. Epilepsy Res Treat. 2012;2012:630853. doi:10.1155/2012/630853

3. Hatton SN, Huynh KH, Bonilha L, et al. White matter abnormalities across different epilepsy syndromes in adults: an ENIGMA-Epilepsy study. Brain. Aug 1 2020;143(8):2454–2473. doi:10.1093/brain/awaa200

4. Whelan CD, Altmann A, Botía JA, et al. Structural brain abnormalities in the common epilepsies assessed in a worldwide ENIGMA study. Brain. Feb 1 2018;141(2):391–408. doi:10.1093/brain/awx341

5. Kandel A, Buzsáki G. Cerebellar neuronal activity correlates with spike and wave EEG patterns in the rat. Epilepsy Res. Sep 1993;16(1):1–9. doi:10.1016/0920-1211(93)90033-4

6. Kros L, Lindeman S, Eelkman Rooda OHJ, et al. Synchronicity and Rhythmicity of Purkinje Cell Firing during Generalized Spike-and-Wave Discharges in a Natural Mouse Model of Absence Epilepsy. Front Cell Neurosci. 2017;11:346. doi:10.3389/fncel.2017.00346

7. Krook-Magnuson E, Szabo GG, Armstrong C, Oijala M, Soltesz I. Cerebellar Directed Optogenetic Intervention Inhibits Spontaneous Hippocampal Seizures in a Mouse Model of Temporal Lobe Epilepsy. eNeuro. Dec 2014;1(1)doi:10.1523/eneuro.0005-14.2014

8. Seto H, Shimizu M, Watanabe N, et al. Contralateral cerebellar activation in frontal lobe epilepsy detected by ictal Tc-99m HMPAO brain SPECT. Clin Nucl Med. Mar 1997;22(3):194–5. doi:10.1097/00003072-199703000-00018

9. Bohnen NI, O’Brien TJ, Mullan BP, So EL. Cerebellar changes in partial seizures: clinical correlations of quantitative SPECT and MRI analysis. Epilepsia. Jun 1998;39(6):640–50. doi:10.1111/j.1528-1157.1998.tb01433.x

10. Marcián V, Mareček R, Koriťáková E, Pail M, Bareš M, Brázdil M. Morphological changes of cerebellar substructures in temporal lobe epilepsy: A complex phenomenon, not mere atrophy. Seizure. Jan 2018;54:51–57. doi:10.1016/j.seizure.2017.12.004

11. Sandok EK, O’Brien TJ, Jack CR, So EL. Significance of cerebellar atrophy in intractable temporal lobe epilepsy: a quantitative MRI study. Epilepsia. Oct 2000;41(10):1315–20. doi:10.1111/j.1528-1157.2000.tb04611.x

12. De Marcos FA, Ghizoni E, Kobayashi E, Li LM, Cendes F. Cerebellar volume and long-term use of phenytoin. Seizure. Jul 2003;12(5):312–5. doi:10.1016/s1059-1311(02)00267-4

13. Hermann BP, Bayless K, Hansen R, Parrish J, Seidenberg M. Cerebellar atrophy in temporal lobe epilepsy. Epilepsy Behav. Sep 2005;7(2):279–87. doi:10.1016/j.yebeh.2005.05.022

14. McDonald CR, Hagler DJ, Jr., Ahmadi ME, et al. Subcortical and cerebellar atrophy in mesial temporal lobe epilepsy revealed by automatic segmentation. Epilepsy Research. May 2008;79(2-3):130–8. doi:10.1016/j.eplepsyres.2008.01.006

15. Oyegbile TO, Bayless K, Dabbs K, et al. The nature and extent of cerebellar atrophy in chronic temporal lobe epilepsy. Epilepsia. Apr 2011;52(4):698–706. doi:10.1111/j.1528-1167.2010.02937.x

16. Hagemann G, Lemieux L, Free SL, et al. Cerebellar volumes in newly diagnosed and chronic epilepsy. J Neurol. Dec 2002;249(12):1651–8. doi:10.1007/s00415-002-0843-9

17. Marcián V, Filip P, Bareš M, Brázdil M. Cerebellar Dysfunction and Ataxia in Patients with Epilepsy: Coincidence, Consequence, or Cause? Tremor Other Hyperkinet Mov (N Y). 2016;6:376. doi:10.7916/d8kh0nbt

18. Bonilha L, Rorden C, Castellano G, et al. Voxel-based morphometry reveals gray matter network atrophy in refractory medial temporal lobe epilepsy. Arch Neurol. Sep 2004;61(9):1379–84. doi:10.1001/archneur.61.9.1379

19. Szabó CA, Lancaster JL, Lee S, et al. MR imaging volumetry of subcortical structures and cerebellar hemispheres in temporal lobe epilepsy. AJNR Am J Neuroradiol. Nov-Dec 2006;27(10):2155–60.

20. Ibdali M, Hadjivassiliou M, Grünewald RA, Shanmugarajah PD. Cerebellar Degeneration in Epilepsy: A Systematic Review. Int J Environ Res Public Health. Jan 8 2021;18(2)doi:10.3390/ijerph18020473

21. Keller SS, Wieshmann UC, Mackay CE, Denby CE, Webb J, Roberts N. Voxel based morphometry of grey matter abnormalities in patients with medically intractable temporal lobe epilepsy: effects of side of seizure onset and epilepsy duration. J Neurol Neurosurg Psychiatry. Dec 2002;73(6):648–55. doi:10.1136/jnnp.73.6.648

22. Keller SS, Wilke M, Wieshmann UC, Sluming VA, Roberts N. Comparison of standard and optimized voxel-based morphometry for analysis of brain changes associated with temporal lobe epilepsy. Neuroimage. Nov 2004;23(3):860–8. doi:10.1016/j.neuroimage.2004.07.030

23. Park KM, Han YH, Kim TH, et al. Cerebellar white matter changes in patients with newly diagnosed partial epilepsy of unknown etiology. Clin Neurol Neurosurg. Nov 2015;138:25–30. doi:10.1016/j.clineuro.2015.07.017

24. Riley JD, Franklin DL, Choi V, et al. Altered white matter integrity in temporal lobe epilepsy: association with cognitive and clinical profiles. Epilepsia. Apr 2010;51(4):536–45. doi:10.1111/j.1528-1167.2009.02508.x

25. Larsell O. The development of the cerebellum in man in relation to its comparative anatomy. The Journal of Comparative Neurology. Oct 1947;87(2):85–129. doi:10.1002/cne.900870203

26. Carass A, Cuzzocreo JL, Han S, et al. Comparing fully automated state-of-the-art cerebellum parcellation from magnetic resonance images. Neuroimage. Dec 2018;183:150–172. doi:10.1016/j.neuroimage.2018.08.003

27. Han S, Carass A, He Y, Prince JL. Automatic cerebellum anatomical parcellation using U-Net with locally constrained optimization. Neuroimage. Sep 2020;218:116819. doi:10.1016/j.neuroimage.2020.116819

28. Ney GC, Lantos G, Barr WB, Schaul N. Cerebellar atrophy in patients with long-term phenytoin exposure and epilepsy. Arch Neurol. Aug 1994;51(8):767–71. doi:10.1001/archneur.1994.00540200043014

29. Blümcke I, Thom M, Aronica E, et al. International consensus classification of hippocampal sclerosis in temporal lobe epilepsy: a Task Force report from the ILAE Commission on Diagnostic Methods. Epilepsia. Jul 2013;54(7):1315–29. doi:10.1111/epi.12220

30. Scheffer IE, Berkovic S, Capovilla G, et al. ILAE classification of the epilepsies: Position paper of the ILAE Commission for Classification and Terminology. Epilepsia. Apr 2017;58(4):512–521. doi:10.1111/epi.13709

31. Kerestes R, Han S, Balachander S, et al. A Standardized Pipeline for Examining Human Cerebellar Grey Matter Morphometry using Structural Magnetic Resonance Imaging. J Vis Exp. Feb 4 2022;(180)doi:10.3791/63340

32. R: A Language and Environment for Statistical Computing. R Foundation for Statistical Computing, Vienna, Austria. 2023.

33. Raz N, Lindenberger U, Rodrigue KM, et al. Regional brain changes in aging healthy adults: general trends, individual differences and modifiers. Cereb Cortex. Nov 2005;15(11):1676–89. doi:10.1093/cercor/bhi044

34. Satterthwaite FE. An approximate distribution of estimates of variance components. Biometrics. Dec 1946;2(6):110–4.

35. Schmahmann JD. An emerging concept. The cerebellar contribution to higher function. Archives of Neurology. Nov 1991;48(11):1178–87. doi:10.1001/archneur.1991.00530230086029

36. Schmahmann JD, Pandya DN. Anatomical investigation of projections to the basis pontis from posterior parietal association cortices in rhesus monkey. The Journal of Comparative Neurology. Nov 1 1989;289(1):53–73. doi:10.1002/cne.902890105

37. Buckner RL, Krienen FM, Castellanos A, Diaz JC, Yeo BT. The organization of the human cerebellum estimated by intrinsic functional connectivity. Journal of Neurophysiology. Nov 2011;106(5):2322–45. doi:10.1152/jn.00339.2011

38. King M, Hernandez-Castillo CR, Poldrack RA, Ivry RB, Diedrichsen J. Functional boundaries in the human cerebellum revealed by a multi-domain task battery. Nature Neuroscience. Aug 2019;22(8):1371–1378. doi:10.1038/s41593-019-0436-x

39. Guo CC, Tan R, Hodges JR, Hu X, Sami S, Hornberger M. Network-selective vulnerability of the human cerebellum to Alzheimer’s disease and frontotemporal dementia. Brain. May 2016;139(Pt 5):1527–38. doi:10.1093/brain/aww003

40. Liang KJ, Carlson ES. Resistance, vulnerability and resilience: A review of the cognitive cerebellum in aging and neurodegenerative diseases. Neurobiol Learn Mem. Apr 2020;170:106981. doi:10.1016/j.nlm.2019.01.004

41. Coan AC, Cendes F. Understanding the spectrum of temporal lobe epilepsy: contributions for the development of individualized therapies. Expert Rev Neurother. Dec 2013;13(12):1383–94. doi:10.1586/14737175.2013.857604

42. Bernhardt BC, Fadaie F, Liu M, et al. Temporal lobe epilepsy: Hippocampal pathology modulates connectome topology and controllability. Neurology. May 7 2019;92(19):e2209–e2220. doi:10.1212/wnl.0000000000007447

43. Bernhardt BC, Bernasconi A, Liu M, et al. The spectrum of structural and functional imaging abnormalities in temporal lobe epilepsy. Ann Neurol. Jul 2016;80(1):142–53. doi:10.1002/ana.24691

44. Liu M, Concha L, Lebel C, Beaulieu C, Gross DW. Mesial temporal sclerosis is linked with more widespread white matter changes in temporal lobe epilepsy. Neuroimage Clin. 2012;1(1):99–105. doi:10.1016/j.nicl.2012.09.010

45. French JA, Bebin M, Dichter MA, et al. Antiepileptogenesis and disease modification: Clinical and regulatory issues. Epilepsia Open. Sep 2021;6(3):483–492. doi:10.1002/epi4.12526

46. Phuong TH, Houot M, Méré M, Denos M, Samson S, Dupont S. Cognitive impairment in temporal lobe epilepsy: contributions of lesion, localization and lateralization. J Neurol. Apr 2021;268(4):1443–1452. doi:10.1007/s00415-020-10307-6

