## Supplementary material for "Patterns of subregional cerebellar atrophy across epilepsy syndromes: An ENIGMA-Epilepsy study"

**Table S1. Demographic and clinical characteristics of epilepsy and control samples, by site**

|  | **Age controls** |  | **Age cases** |  | **Female % Age of onset** | | | | |  | **Duration epilepsy** |  | **Total control** |  | **Total cases** |  | | **TLE-HS-L** | |  | | **TLE-HS-R** | |  | | **TLE-NL-L** | |  | | **TLE-NL-R** | |  | | **GGE** | |  | | **ETLE** | |
| --- | --- | --- | --- | --- | --- | --- | --- | --- | --- | --- | --- | --- | --- | --- | --- | --- | --- | --- | --- | --- | --- | --- | --- | --- | --- | --- | --- | --- | --- | --- | --- | --- | --- | --- | --- | --- | --- | --- | --- |
| **Site** | **Mean (SD)** |  | **Mean (SD)** |  | **Controls** |  | **Cases** |  | **Mean (SD)** |  |  |  |  |  |  |  | |  | |  | |  | |  | |  | |  | |  | |  | |  | |  | |  | |
| Bern | 32.5 (9.39) |  | 33.6 (11.9) |  | 53 |  | 55 |  | - |  | - |  | 78 |  | 109 |  | | 18 | |  | | 13 | |  | | 18 | |  | | 13 | |  | | 11 | |  | | 33 | |
| Bonn |  |  | 38.3 (11.2) |  |  |  | 52 |  | 20.3 (16.5) |  | 15.6 (15.1) |  | 0 |  | 259 |  | | 66 | |  | | 43 | |  | | 30 | |  | | 16 | |  | | 0 | |  | | 0 | |
| CapeTown | 52.5 (7.78) |  | 37.5 (11.4) |  | 50 |  | 56 |  | 29.2 (14.8)** |  | 8.1 (6.4)** |  | 2 |  | 39 |  | | 5 | |  | | 3 | |  | | 5 | |  | | 12 | |  | | 0 | |  | | 4 | |
| CUBRIC | 27.4 (7.82) |  | 27.5 (7.4) |  | 74 |  | 73 |  | 12.3 (4.5) |  | 15.2 (9.1) |  | 42 |  | 40 |  | | 0 | |  | | 0 | |  | | 0 | |  | | 0 | |  | | 40 | |  | | 0 | |
| EKUT_Skyra | 34.1 (11.20) |  | 36.2 (13.0) |  | 55 |  | 50 |  | 18.7 (12.9)* |  | 18.4 (13.8)* |  | 78 |  | 121 |  | | 15 | |  | | 13 | |  | | 12 | |  | | 4 | |  | | 4 | |  | | 67 | |
| EKUT_Prisma | 29.9 (8.90) |  | 27.3 (9.6) |  | 49 |  | 42 |  | 14.3 (9.5)* |  | 12.8 (9.89)* |  | 85 |  | 48 |  | | 0 | |  | | 1 | |  | | 0 | |  | | 2 | |  | | 20 | |  | | 15 | |
| EPICZ | 38.5 (10.77) |  | 38.8 (9.5) |  | 50 |  | 64 |  | 20.7 (11.9) |  | 18.22 (12.2) |  | 109 |  | 105 |  | | 20 | |  | | 16 | |  | | 35 | |  | | 28 | |  | | 0 | |  | | 4 | |
| Florence | 35.3 (8.48) |  | 27.8 (7.8) |  | 57 |  | 40 |  | 12.8 (8.2)* |  | 14.3 (8.0)* |  | 14 |  | 30 |  | | 1 | |  | | 1 | |  | | 1 | |  | | 0 | |  | | 5 | |  | | 19 | |
| Genova | 28.2 (11.07) |  | 23.6 (7.7) |  | 68 |  | 65 |  | 11.6 (7.2) |  | 12.0 (4.8) |  | 31 |  | 20 |  | | 0 | |  | | 1 | |  | | 5 | |  | | 2 | |  | | 7 | |  | | 3 | |
| IDIBAPS | 33.3 (7.37) |  | 35.9 (10.1) |  | 62 |  | 62 |  | 15.2 (10.3)* |  | 18.5 (11.7)* |  | 68 |  | 251 |  | | 45 | |  | | 45 | |  | | 15 | |  | | 19 | |  | | 11 | |  | | 61 | |
| RMH |  |  | 36.6 (12.9) |  |  |  | 49 |  | 27.1 (15.1) |  | 9.7 (12.5) |  | 0 |  | 158 |  | | 16 | |  | | 10 | |  | | 8 | |  | | 10 | |  | | 5 | |  | | 13 | |
| UCL | 37.6 (12.40) |  | 38.0 (10.9) |  | 58 |  | 62 |  | 12.9 (9.6) |  | 25.1 (14.5) |  | 29 |  | 50 |  | | 24 | |  | | 12 | |  | | 5 | |  | | 9 | |  | | 0 | |  | | 0 | |
| UCLA | 39.23 (9.21) |  | 36.1 (10.4) |  | 64 |  | 55 |  | 22.9 (12.1) |  | 13.2 (10.5) |  | 45 |  | 20 |  | | 1 | |  | | 1 | |  | | 5 | |  | | 1 | |  | | 0 | |  | | 5 | |
| UCSD | 38.0 (14.50) |  | 34.34 (10.5) |  | 41 |  | 50 |  | 19.6 (13.3) |  | 14.7 (11.2) |  | 22 |  | 50 |  | | 12 | |  | | 9 | |  | | 11 | |  | | 13 | |  | | 0 | |  | | 0 | |
| UNICAMP | 34.34 (10.46) |  | 41.9 (9.2) |  | 62 |  | 62 |  | 12.0 (9.3) |  | 29.9 (12.6) |  | 382 |  | 209 |  | | 92 | |  | | 72 | |  | | 0 | |  | | 0 | |  | | 34 | |  | | 0 | |
| UNIMORE | 28.2 (5.12) |  | 30.1 (10.4) |  | 54 |  | 58 |  | 15.4 (10.9)* |  | 14.5 (11.6)* |  | 37 |  | 93 |  | | 5 | |  | | 2 | |  | | 3 | |  | | 2 | |  | | 49 | |  | | 27 | |
| **Total** |  |  |  |  |  |  |  |  |  |  |  |  | **1022** |  | **1602** | |  | | **320** | |  | | **242** | |  | | **153** | |  | | **131** | |  | | **186** | |  | | **251** |

**Table S2**. Breakdown of diagnoses in the 'all other epilepsies' aggregate group

|  | **N** |  | **% of sample** |
| --- | --- | --- | --- |
| **Focal- temporal lobe epilepsies** |  |  |  |
| Bilateral TLE with HS | 7 |  | 2.2 |
| Bilateral TLE without HS (lesion-negative) | 11 |  | 3.5 |
| Bilateral TLE, information on HS not available | 40 |  | 12.6 |
| Unclear seizure focus | 189 |  | 59.6 |
| Unknown/inconclusive | 67 |  | 21.1 |
| Other | 3 |  | 1 |
| **Total** | **317** |  |  |

**Table S3.** Scanner protocols by each research site.

| **Site** | **Scanner Brand/Model** | **Field Strength (Tesla)** | **Head Coil (Channels)** | **Sequence Name** | **TR (ms)** | **TE (ms)** | **TI (ms)** | **Flip Angle** | **Acquisition Name** | **# of Slices** | **Field of View** | **Voxel Size (mm3)** |
| --- | --- | --- | --- | --- | --- | --- | --- | --- | --- | --- | --- | --- |
| **Bern_1** | Siemens Trio | 3 |  | MPRAGE | 1860 | 3.42 | 800 | 15 | Sagittal | 160 | 256 x 256 | 1 x 1 x 1 |
| **Bern_2** | Siemens Trio Tim | 3 |  | MDEFT3d1 | 7.92 | 2.48 | 910 | 16 | Sagittal | 176 | 256 x 256 | 1 x 1 x 1 |
| **Bonn_1** | Siemens Trio | 3 | 32 | t1_mpr_32ch_iso_0.8 | 1660 | 2.54 | 850 | 9 | Sagittal | 208 | 256 x 256 | 0.8 x 0.8 x 0.8 |
| **Bonn_2** | Siemens Trio | 3 | 8 | t1_mpr_ns_sag_1x1x1 | 1570 | 3.42 | 800 | 15 | Sagittal | 160 | 256 x 256 | 1 x 1 x 1 |
| **Bonn_3** | Philips Achieva TX | 3 | 8 | T1_Navigato | 7.53 | 3.48 | - | 8 | Sagittal | 170 | 256 x 256 | 1 x 1 x 1 |
| **Bonn_4** | Philips Intera | 3 | - | S3DTFESF3.6 SENSE | 8.10 | 3.70 | - | 8 | Sagittal | 140 | 256 x 256 | 1 x 1 x 1 |
| **CUBRIC** | GESigna HDx | 3 | 8 | FSPGR | 8.00 | 3.00 | 450 | 20 | Axial | - | 256 x 256 | 1 x 1 x 1 |
| **EKUT_Prisma** | Siemens Prisma | 3 | 64 | MPRAGE | 2300 | 2.98 | 900 | 9 | Axial | 256 | 240 x 256 | 1 x 1 x 1 |
| **EKUT_Skyra** | Siemens Skyra | 3 | 32 | MPRAGE | 2300 | 2.32 | 900 | 8 | Axial | 256 | 172 x 230 | 0.9 x 0.9 x 0.9 |
| **EPICZ** | GE Discovery | 3 | 8 | BRAVO | 9.20 | 3.70 | - | 12 | Sagittal | 368 | 256 x 256 | 1 x 1 x 1 |
| **Florence_1** | Philips Achieva | 3 | 32 | 3D T1-weighted turbo field echo | 8.10 | 3.70 | - | 8 | Sagittal | 191 | 256 x 256 | 1 x 1 x 1 |
| **Florence_2** | Philips Achieva | 1.5 | 32 | 3D T1-weighted turbo field echo | 25 | 4.60 | - | 30 | Sagittal | 175 | 256 x 256 | 1 x 1 x 1 |
| **IDIBAPS** | Siemens Trio Tim | 3 | 32 | CORONAL 3D MPRAGE | 2000 | 3.03 | 900 | 9 | Coronal | 192 | 256 x 224 | 0.9 x 1.2 x 0.9 |
| **IGG_1** | Philips | 3 | 32 | T1 vol neuronavigazione | 7.90 | 3.50 | 0 | 8 | Axial | 160 | 256 x 256 | 1 x 1 x 1 |
| **IGG_2** | Philips | 1.5 | 8 | sT1W_3D_TFE SENSE | 7.04 | 3.19 | 0 | 8 | Sagittal | 162 | 180 x 180 | 1 x 1 x 1 |
| **RMH** | Siemens | 3 | 32 | tfl3d1_ns | 1900 | 2.21 | 900 | 9 | Coronal | 192 | - | 0.5 x 0.5 x 1 |
| **UCL** | GE | 3 | 8 | FSPGR 3D | 8.00 | 3.00 | 450 | 20 | Coronal | 170 | 256 x 256 | 1 x 1 x 1 |
| **UCLA** | Siemens Skyra | 3 | 20 | MPRAGE | 11.00 | 2.81 | 1100 | 20 | Axial | 176 | 256 x 256 | 1 x 1 x 1 |
| **UCSD** | GE Discovery | 3 | 8 | FSPGR_SAG_TI450 | 8.13 | 3.17 | 600 | 8 | Axial | 172 | 256 x 256 | 1 x 1 x 1 |
| **UCT_Capetown** | Avanto MRI | 1.5 | 20 | t1_mprage-sag | 2400 | 3.61 | - | 8 | Sagittal | 160 | 240 x 240 | 1 x 1 x 1 |
| **UNICAMP** | Philips Achieva | 3 | 8 | - | 7.00 | 3.20 | - | 8 | Sagittal | 180 | 240 x 240 | 1 x 1 x 1 |
| **UNIMORE** | Philips Ingenia | 3 | 32 | T1-3D MPRAGE | 9.90 | 4.60 | 670 | 8 | Sagittal | 170 | 210 x 240 | 1 x 1 x 1 |

**Laterality analyses**

We first summed all left cerebellar ROI volumes and all right cerebellar ROI volumes (excluding vermis and corpus medullare), separately for each person in the TLE-HS-L, TLE-HS-R, TLE- L-NL and TLE-R-NL groups. Given the moderate-high degree of symmetry in cerebellar volumes between hemispheres across all participants (*r*>0.7), outlier volumes (i.e., volumes subsequently treated as NA) were first imputed based on all other volumes including total volume and ICV using the R package *missRanger.* A laterality index was defined as the relative proportion of volume in the right cerebellar hemisphere relative to total cerebellar volume (i.e., 0.5 = no laterality; >0.5 = more volume on right; <0.5 = more volume on left). A linear model was fit to the data, with laterality as a predictor variable.

**Table S4:** Effect sizes, betas, standard error and FDR-adjusted P values for cerebellum volume differences between all epilepsies and healthy controls controlling for age, age^2^, sex, ICV and site

| Cerebellar region | Cohen's *d* | Lower CI | Upper CI | Beta | Std error | Statistic (*t*) | FDR-adjusted *P* value |
| --- | --- | --- | --- | --- | --- | --- | --- |
| Corpus.Medullare | 0.4874369 | 0.3908643 | 0.5840095 | -1017.8902 | 96.351256 | -10.5643685 | **0** |
| Left.VIIB | 0.4015032 | 0.3061391 | 0.4968672 | -437.33607 | 50.044418 | -8.7389579 | **0** |
| Right.VIIB | 0.3250698 | 0.2305602 | 0.4195795 | -391.49192 | 55.095729 | -7.1056672 | **0** |
| Right.Crus.II | 0.3152441 | 0.2203122 | 0.4101759 | -424.01332 | 61.618549 | -6.881261 | **0** |
| Right.V | 0.2694746 | 0.1747398 | 0.3642093 | -125.90082 | 21.491143 | -5.8582655 | **0** |
| Left.Crus.II | 0.2649625 | 0.1701606 | 0.3597644 | -349.97025 | 60.466781 | -5.7878101 | **0** |
| Left.VIIIB | 0.2541731 | 0.159524 | 0.3488221 | -164.69921 | 29.825769 | -5.5220441 | **0.0000001** |
| Left.Crus.I | 0.2468354 | 0.1532178 | 0.3404529 | -469.28632 | 86.759046 | -5.4090765 | **0.0000002** |
| Left.IX | 0.2462414 | 0.1514675 | 0.3410153 | -160.76227 | 29.84328 | -5.3868833 | **0.0000002** |
| Right.VIIIB | 0.2454389 | 0.149699 | 0.3411788 | -160.43777 | 30.270508 | -5.3001347 | **0.0000004** |
| Vermis.IX | 0.2364732 | 0.1423936 | 0.3305528 | -36.158408 | 6.982344 | -5.1785489 | **0.0000006** |
| Right.IX | 0.2272378 | 0.1336003 | 0.3208753 | -147.95911 | 29.6731 | -4.9863045 | **0.0000015** |
| Left.VIIIA | 0.2172201 | 0.1220065 | 0.3124337 | -237.86773 | 50.387151 | -4.7208014 | **0.0000053** |
| Right.Crus.I | 0.2084642 | 0.1147016 | 0.3022268 | -389.3755 | 85.339737 | -4.5626518 | **0.0000106** |
| Left.VI | 0.1825202 | 0.0896975 | 0.2753429 | -236.15429 | 58.887075 | -4.0102907 | **0.0001166** |
| Vermis.X | 0.1787233 | 0.0849314 | 0.2725151 | -10.300746 | 2.628765 | -3.9184738 | **0.0001601** |
| Vermis.VIII | 0.1568713 | 0.0628331 | 0.2509094 | -46.030057 | 13.426438 | -3.4283149 | **0.0010161** |
| Vermis.VII | 0.1399835 | 0.0460115 | 0.2339555 | -24.032074 | 7.849544 | -3.0615885 | **0.0034606** |
| Right.VIIIA | 0.1230763 | 0.0277409 | 0.2184116 | -117.98957 | 44.32565 | -2.6618802 | **0.0114674** |
| Right.X | 0.12137 | 0.0267389 | 0.2160012 | -10.118093 | 3.823714 | -2.6461426 | **0.0114674** |
| Right.I.III | 0.1170321 | 0.0233194 | 0.2107448 | -21.976017 | 8.579012 | -2.5616021 | **0.0139686** |
| Left.V | 0.1014756 | 0.0067445 | 0.1962067 | -47.773863 | 21.662949 | -2.2053259 | **0.0350235** |
| Left.I.III | 0.0849466 | -0.0084789 | 0.1783721 | -14.899891 | 7.990077 | -1.8647995 | 0.0741875 |
| Left.IV | -0.0852287 | -0.1798227 | 0.0093653 | 46.8727414 | 25.256852 | 1.8558426 | 0.0741875 |
| Right.VI | 0.0568258 | -0.0367368 | 0.1503883 | -71.84755 | 57.802322 | -1.2429873 | 0.2396652 |
| Left.X | 0.0465675 | -0.0480375 | 0.1411725 | -3.962373 | 3.899249 | -1.0161886 | 0.333454 |
| Right.IV | 0.0131753 | -0.081948 | 0.1082987 | -7.3797832 | 25.7984 | -0.2860558 | 0.8035565 |
| Vermis.VI | -0.0008399 | -0.0937479 | 0.092068 | 0.1942262 | 10.535113 | 0.0184361 | 0.9852925 |

**Table S5.** Effect sizes, betas, standard error and FDR-adjusted P values for cerebellum volume differences between TLE-HS and healthy controls, controlling for age, age^2^, sex, ICV and site

| Cerebellar region | Cohen's *d* | Lower CI | Upper CI | Beta | Std error | Statistic (*t*) | FDR-adjusted *P* value |
| --- | --- | --- | --- | --- | --- | --- | --- |
| **TLE-HS-L** |  |  |  |  |  |  |  |
| Corpus.Medullare | 0.5662362 | 0.4049655 | 0.7275069 | -1103.155 | 150.347089 | -7.3373883 | **0** |
| Left.VIIB | 0.476302 | 0.3184103 | 0.6341936 | -514.06821 | 81.885976 | -6.2778541 | **0** |
| Right.VIIB | 0.4551293 | 0.3036984 | 0.6065603 | -542.27409 | 87.519505 | -6.196037 | **0** |
| Right.V | 0.4102454 | 0.2534956 | 0.5669953 | -191.87186 | 35.537425 | -5.3991493 | **0.0000006** |
| Right.Crus.II | 0.3979034 | 0.240204 | 0.5556027 | -519.46982 | 97.811727 | -5.3109156 | **0.0000008** |
| Vermis.IX | 0.3440189 | 0.1887511 | 0.4992866 | -52.023733 | 11.236395 | -4.6299308 | **0.0000195** |
| Left.Crus.I | 0.3308949 | 0.1771584 | 0.4846314 | -610.83307 | 137.227324 | -4.4512496 | **0.0000382** |
| Right.VIIIB | 0.3182582 | 0.1604067 | 0.4761098 | -201.15123 | 48.128595 | -4.1794535 | **0.0001007** |
| Left.Crus.II | 0.3139844 | 0.1579239 | 0.4700448 | -407.84674 | 97.707524 | -4.174159 | **0.0001007** |
| Left.VIIIB | 0.3012644 | 0.1462752 | 0.4562536 | -191.51126 | 47.814338 | -4.0053102 | **0.0001852** |
| Right.IX | 0.2846017 | 0.1342421 | 0.4349612 | -182.37391 | 46.92558 | -3.8864498 | **0.0002794** |
| Left.VIIIA | 0.2869771 | 0.1311662 | 0.4427881 | -304.30469 | 80.062476 | -3.8008404 | **0.0003539** |
| Right.Crus.I | 0.2813134 | 0.126842 | 0.4357849 | -501.23523 | 133.169701 | -3.7638834 | **0.0003809** |
| Left.VI | 0.2755048 | 0.1237927 | 0.4272169 | -349.57534 | 93.686013 | -3.7313504 | **0.000404** |
| Left.IX | 0.2744599 | 0.1181572 | 0.4307625 | -172.93318 | 47.177639 | -3.6655751 | **0.0004842** |
| Vermis.X | 0.2533405 | 0.0987318 | 0.4079492 | -13.863217 | 4.064766 | -3.410582 | **0.001183** |
| Left.V | 0.2336966 | 0.0759352 | 0.3914579 | -106.75509 | 34.979597 | -3.0519246 | **0.0036845** |
| Right.X | 0.2301427 | 0.0734155 | 0.38687 | -18.9529 | 6.221432 | -3.0463885 | **0.0036845** |
| Vermis.VIII | 0.2218773 | 0.0666612 | 0.3770935 | -63.441642 | 21.527431 | -2.947014 | **0.0048238** |
| Vermis.VII | 0.1728122 | 0.022419 | 0.3232054 | -29.196594 | 12.446955 | -2.3456816 | **0.0268883** |
| Left.IV | -0.1647032 | -0.3213008 | -0.0081057 | 89.3544175 | 41.169256 | 2.1704161 | **0.0402262** |
| Left.I.III | 0.1552716 | 0.002304 | 0.3082393 | -27.033602 | 12.970565 | -2.0842271 | **0.0475917** |
| Right.I.III | 0.1529363 | -0.0005315 | 0.3064041 | -28.217857 | 13.801055 | -2.0446159 | 0.0500769 |
| Right.VIIIA | 0.1292906 | -0.025347 | 0.2839281 | -125.56604 | 73.08515 | -1.7180786 | 0.0966772 |
| Left.X | 0.1291327 | -0.0270825 | 0.2853479 | -10.488205 | 6.109499 | -1.7167047 | 0.0966772 |
| Right.VI | 0.1266888 | -0.0263861 | 0.2797638 | -159.85849 | 94.203512 | -1.6969483 | 0.0969213 |
| Vermis.VI | -0.0036561 | -0.1536221 | 0.1463099 | 0.8440435 | 17.022125 | 0.0495851 | 0.9604635 |
| **TLE-HS-R** |  |  |  |  |  |  |  |
| Corpus.Medullare | 0.6273514 | 0.4507242 | 0.8039787 | -1224.761 | 164.923647 | -7.4262301 | **0** |
| Left.VIIB | 0.421959 | 0.2506517 | 0.5932663 | -446.18958 | 87.675618 | -5.0890953 | **0.0000059** |
| Right.VIIB | 0.3783908 | 0.2126126 | 0.544169 | -450.88341 | 96.188708 | -4.687488 | **0.0000296** |
| Right.V | 0.340394 | 0.1690243 | 0.5117637 | -161.2884 | 39.377067 | -4.0959982 | **0.000315** |
| Left.Crus.I | 0.3202809 | 0.1575873 | 0.4829745 | -592.19366 | 147.568804 | -4.0130003 | **0.0003488** |
| Left.Crus.II | 0.3264972 | 0.1568031 | 0.4961913 | -416.19596 | 104.659512 | -3.9766663 | **0.0003488** |
| Vermis.IX | 0.3169286 | 0.1465876 | 0.4872696 | -47.630523 | 12.27177 | -3.8813083 | **0.0004446** |
| Right.Crus.II | 0.297498 | 0.1279667 | 0.4670293 | -383.99869 | 104.696752 | -3.6677231 | **0.0009066** |
| Left.IX | 0.2814963 | 0.1115366 | 0.4514559 | -179.01989 | 51.719155 | -3.4613846 | **0.0017474** |
| Left.VIIIA | 0.247256 | 0.0748393 | 0.4196728 | -258.22957 | 87.042458 | -2.9667081 | **0.0086003** |
| Right Crus I | 0.2328326 | 0.0672367 | 0.3984285 | -418.59872 | 145.264338 | -2.8816344 | **0.0102977** |
| Right.X | 0.2285208 | 0.0560342 | 0.4010075 | -18.975422 | 6.879047 | -2.7584375 | **0.0128203** |
| Right.IX | 0.2199427 | 0.0560522 | 0.3838333 | -142.71184 | 51.757947 | -2.7572934 | **0.0128203** |
| Left.VI | 0.207817 | 0.0448386 | 0.3707953 | -265.50548 | 101.843288 | -2.6070003 | **0.0185814** |
| Vermis.VIII | 0.1957615 | 0.027014 | 0.364509 | -56.505941 | 23.71239 | -2.3829712 | **0.0323835** |
| Vermis.X | 0.171489 | 0.0067099 | 0.3362681 | -9.576831 | 4.436615 | -2.1585896 | 0.0545843 |
| Vermis.VII | 0.1610751 | -0.006292 | 0.3284422 | -26.781731 | 13.564159 | -1.9744483 | 0.0800502 |
| Left.X | 0.1573277 | -0.0158515 | 0.330507 | -13.382599 | 7.041172 | -1.9006209 | 0.0896268 |
| Left.IV | -0.1444907 | -0.3141564 | 0.025175 | 78.135028 | 44.723732 | 1.7470596 | 0.1192156 |
| Right.VIIIB | 0.139502 | -0.0338092 | 0.3128131 | -86.184285 | 51.70472 | -1.6668553 | 0.1341385 |
| Right.VIIIA | 0.1321629 | -0.0395264 | 0.3038522 | -125.91059 | 79.155823 | -1.5906674 | 0.1492915 |
| Right.I.III | 0.1114062 | -0.0561391 | 0.2789514 | -21.168797 | 15.553334 | -1.3610456 | 0.2211851 |
| Left.VIIIB | 0.1052934 | -0.0649619 | 0.2755487 | -66.586758 | 52.153617 | -1.2767429 | 0.2361885 |
| Vermis.VI | 0.1033469 | -0.0627716 | 0.2694653 | -23.146655 | 18.256608 | -1.2678508 | 0.2361885 |
| Right.VI | 0.1016878 | -0.0650091 | 0.2683848 | -129.35864 | 103.324651 | -1.251963 | 0.2361885 |
| Right.IV | -0.0767514 | -0.2483793 | 0.0948765 | 41.107247 | 44.550799 | 0.922705 | 0.3837625 |
| Left.I.III | 0.063058 | -0.104923 | 0.231039 | -11.249796 | 14.567425 | -0.772257 | 0.4564437 |
| Left.V | -0.0087691 | -0.1809684 | 0.1634303 | 4.026048 | 38.425596 | 0.1047752 | 0.9165716 |

**Table S6.** Effect sizes, betas, standard error and FDR-adjusted P values for cerebellum volume differences between TLE-NL and healthy controls, controlling for age, age^2^, sex, ICV and site

| Cerebellar region | Cohen's *d* | Lower CI | Upper CI | Beta | Std error | Statistic (*t*) | FDR-adjusted *P* value |
| --- | --- | --- | --- | --- | --- | --- | --- |
| **TLE-NL-L** |  |  |  |  |  |  |  |
| Corpus.Medullare | 0.5805191 | 0.3622595 | 0.7987786 | -1089.5651 | 196.901472 | -5.5335547 | **0.0000011** |
| Left.X | -0.2520042 | -0.4563013 | -0.047707 | 20.672852 | 8.065804 | 2.5630244 | 0.1483244 |
| Right.V | 0.203683 | -0.0049646 | 0.4123307 | -96.365885 | 48.055661 | -2.0052972 | 0.3247333 |
| Vermis.VIII | 0.1881067 | -0.0188765 | 0.39509 | -53.987464 | 28.852266 | -1.8711689 | 0.3247333 |
| Vermis.VII | 0.1834972 | -0.0190504 | 0.3860447 | -30.500127 | 16.384705 | -1.8614999 | 0.3247333 |
| Vermis.IX | 0.1783463 | -0.0252591 | 0.3819517 | -26.602034 | 14.637438 | -1.8173968 | 0.3247333 |
| Vermis.VI | 0.1614052 | -0.0396724 | 0.3624828 | -36.941408 | 22.553398 | -1.6379531 | 0.4072806 |
| Right.VIIIB | 0.1639468 | -0.054536 | 0.3824296 | -98.057549 | 62.90941 | -1.5587104 | 0.4177988 |
| Left.VIIB | 0.1469564 | -0.062138 | 0.3560508 | -154.17171 | 106.508734 | -1.4475029 | 0.4212679 |
| Right.Crus.II | 0.1380474 | -0.0615151 | 0.33761 | -175.08872 | 121.597602 | -1.4399027 | 0.4212679 |
| Vermis.X | 0.1122885 | -0.0871779 | 0.3117548 | -6.178161 | 5.311921 | -1.1630749 | 0.6050627 |
| Left.VI | 0.107699 | -0.0888986 | 0.3042966 | -136.3342 | 121.874343 | -1.1186456 | 0.6050627 |
| Right.VIIB | -0.1028645 | -0.3034732 | 0.0977442 | 122.314069 | 116.903204 | 1.046285 | 0.6050627 |
| Right.X | 0.1030275 | -0.1036099 | 0.3096649 | -8.532278 | 8.270138 | -1.0316973 | 0.6050627 |
| Left.VIIIB | 0.0778209 | -0.1332405 | 0.2888824 | -48.021477 | 62.959698 | -0.7627336 | 0.8048837 |
| Right.IX | 0.072897 | -0.130974 | 0.2767681 | -46.800421 | 63.297449 | -0.7393729 | 0.8048837 |
| Left.I.III | 0.0646449 | -0.1392476 | 0.2685374 | -11.316762 | 17.421775 | -0.6495757 | 0.8501219 |
| Right.I.III | 0.0534461 | -0.152634 | 0.2595261 | -10.180748 | 19.106622 | -0.5328387 | 0.8663213 |
| Left.IX | 0.0503286 | -0.158428 | 0.2590851 | -31.813049 | 63.244167 | -0.5030195 | 0.8663213 |
| Right.IV | 0.0434479 | -0.1716779 | 0.2585737 | -23.218745 | 55.095234 | -0.4214293 | 0.8663213 |
| Left.Crus.II | 0.0364014 | -0.1707158 | 0.2435186 | -45.505311 | 125.340688 | -0.363053 | 0.8663213 |
| Left.Crus.I | 0.0329654 | -0.1660136 | 0.2319443 | -58.629909 | 173.019258 | -0.3388635 | 0.8663213 |
| Left.V | 0.033278 | -0.1794434 | 0.2459993 | -15.238755 | 47.16726 | -0.3230791 | 0.8663213 |
| Left.IV | -0.0322607 | -0.2438329 | 0.1793116 | 16.887719 | 53.745883 | 0.3142142 | 0.8663213 |
| Right.VIIIA | -0.0265908 | -0.2323312 | 0.1791496 | 25.219663 | 95.108215 | 0.2651681 | 0.8663213 |
| Right.Crus.I | -0.0244031 | -0.2260582 | 0.177252 | 42.546554 | 171.772202 | 0.2476917 | 0.8663213 |
| Right.VI | -0.0127552 | -0.213192 | 0.1876816 | 16.04225 | 123.8715 | 0.1295072 | 0.9236242 |
| **TLE-NL-R** |  |  |  |  |  |  |  |
| Corpus.Medullare | 0.5672897 | 0.3301837 | 0.8043957 | -1087.7669 | 217.181766 | -5.0085553 | **0.0000179** |
| Left.IV | -0.3811548 | -0.608533 | -0.1537766 | 203.862562 | 58.695878 | 3.4732007 | **0.0075474** |
| Vermis.IX | 0.2984285 | 0.07306 | 0.523797 | -44.477342 | 15.99098 | -2.781402 | 0.0518642 |
| Vermis.VIII | 0.2383608 | 0.0148541 | 0.4618676 | -67.751359 | 30.756992 | -2.2027954 | 0.1951904 |
| Right.IV | -0.2247764 | -0.4558407 | 0.0062878 | 122.13091 | 60.096552 | 2.0322449 | 0.2313076 |
| Left.VIIIA | 0.2100784 | -0.0136277 | 0.4337846 | -213.19933 | 110.19353 | -1.9347717 | 0.2313076 |
| Vermis.VI | -0.2003947 | -0.4187488 | 0.0179594 | 45.7162052 | 24.277192 | 1.8830928 | 0.2313076 |
| Left.V | -0.2042602 | -0.4334893 | 0.0249689 | 94.6079315 | 51.420104 | 1.8399016 | 0.2313076 |
| Left.VIIB | 0.1719472 | -0.0516017 | 0.3954961 | -180.18564 | 113.721822 | -1.5844421 | 0.3529985 |
| Right.Crus.II | 0.1509019 | -0.0694198 | 0.3712235 | -189.78452 | 130.199223 | -1.4576471 | 0.3636761 |
| Right.IX | 0.1506237 | -0.0657027 | 0.3669502 | -95.526994 | 66.461117 | -1.4373366 | 0.3636761 |
| Left.VI | 0.1433603 | -0.0698066 | 0.3565272 | -181.61734 | 131.303418 | -1.3831882 | 0.3636761 |
| Left.I.III | -0.1478136 | -0.3688567 | 0.0732296 | 25.9924278 | 18.873568 | 1.3771868 | 0.3636761 |
| Right.X | 0.1336649 | -0.086909 | 0.3542388 | -11.042559 | 8.811479 | -1.2532016 | 0.4210609 |
| Left.Crus.I | 0.123523 | -0.092537 | 0.339583 | -221.36616 | 187.633262 | -1.179781 | 0.4339749 |
| Left.VIIIB | 0.1200987 | -0.1046912 | 0.3448885 | -75.643852 | 68.553446 | -1.1034289 | 0.4339749 |
| Right.VIIIB | 0.1205608 | -0.1091312 | 0.3502527 | -73.044653 | 67.063807 | -1.0891814 | 0.4339749 |
| Left.Crus.II | 0.1112691 | -0.1111318 | 0.33367 | -139.99869 | 135.358951 | -1.0342773 | 0.4339749 |
| Left.IX | 0.1092967 | -0.1122175 | 0.3308108 | -67.562292 | 65.587549 | -1.0301085 | 0.4339749 |
| Vermis.X | 0.1064491 | -0.1123402 | 0.3252384 | -5.7539131 | 5.66265 | -1.0161166 | 0.4339749 |
| Right.Crus.I | 0.0614317 | -0.1590475 | 0.2819108 | -107.28279 | 185.366026 | -0.5787619 | 0.7505965 |
| Right.V | 0.051296 | -0.1748013 | 0.2773933 | -24.624849 | 52.675886 | -0.4674786 | 0.7994656 |
| Right.VIIIA | 0.0480237 | -0.1735725 | 0.2696199 | -45.090153 | 101.408349 | -0.4446395 | 0.7994656 |
| Vermis.VII | -0.0401007 | -0.2567615 | 0.1765601 | 6.8231561 | 17.96798 | 0.3797397 | 0.8216248 |
| Right.I.III | -0.0272671 | -0.2478344 | 0.1933002 | 5.1710826 | 20.390499 | 0.2536026 | 0.8958523 |
| Right.VI | -0.0201094 | -0.2355149 | 0.1952962 | 25.0671157 | 130.824647 | 0.1916085 | 0.9133434 |
| Right.VIIB | 0.0141068 | -0.2021704 | 0.230384 | -16.768754 | 125.020464 | -0.1341281 | 0.9246177 |
| Left.X | 0.0099256 | -0.2078287 | 0.2276798 | -0.8235589 | 8.700439 | -0.0946572 | 0.9246177 |
| Left.V | -0.0087691 | -0.1809684 | 0.1634303 | 4.026048 | 38.425596 | 0.1047752 | 0.9165716 |

**Table S7.** Effect sizes, betas, standard error and FDR-adjusted P values for cerebellum volume differences between GGE and healthy controls, controlling for age, age^2^, sex, ICV and site

| Cerebellar region | Cohen's *d* | Lower CI | Upper CI | Beta | Std error | Statistic (*t*) | FDR-adjusted *P* value |
| --- | --- | --- | --- | --- | --- | --- | --- |
| **GGE** |  |  |  |  |  |  |  |
| Corpus.Medullare | 0.4667068 | 0.2744746 | 0.658939 | -882.30036 | 172.677117 | -5.1095384 | **0.0000105** |
| Left.VIIB | 0.383735 | 0.1957398 | 0.5717301 | -395.91414 | 93.158021 | -4.24992 | **0.0003229** |
| Right.Crus.I | 0.3468863 | 0.1619368 | 0.5318359 | -598.46861 | 153.894476 | -3.8888245 | **0.0009967** |
| Right.VIIB | 0.2859792 | 0.0982888 | 0.4736695 | -338.26018 | 106.651834 | -3.1716302 | **0.0088505** |
| Left.VIIIA | 0.2868497 | 0.0969652 | 0.4767342 | -288.96864 | 91.248828 | -3.1668203 | **0.0088505** |
| Left.Crus.I | 0.272095 | 0.0880018 | 0.4561883 | -484.05506 | 158.113658 | -3.0614374 | **0.0105368** |
| Right.VIIIB | 0.2706985 | 0.0792827 | 0.4621143 | -159.18942 | 53.588225 | -2.9706044 | **0.0110347** |
| Vermis.VIII | 0.2676587 | 0.0788428 | 0.4564745 | -76.146304 | 25.737973 | -2.9585198 | **0.0110347** |
| Right.V | 0.2472407 | 0.0586881 | 0.4357932 | -117.31874 | 42.947594 | -2.7316719 | **0.0198953** |
| Left.VIIIB | 0.2371751 | 0.0494295 | 0.4249208 | -146.63474 | 55.81366 | -2.6272195 | **0.0244222** |
| Left.IX | 0.1995805 | 0.0090823 | 0.3900786 | -124.51398 | 56.084877 | -2.2200989 | 0.0677353 |
| Right.VIIIA | 0.1838144 | -0.0061898 | 0.3738186 | -171.93153 | 85.081794 | -2.0207794 | 0.1014613 |
| Right.IX | 0.1781683 | -0.0094881 | 0.3658247 | -112.49547 | 56.600239 | -1.9875441 | 0.1014613 |
| Left.Crus.II | 0.1596737 | -0.0314909 | 0.3508383 | -204.52143 | 115.628303 | -1.7687835 | 0.1543772 |
| Right.Crus.II | 0.1454371 | -0.042962 | 0.3338361 | -187.10301 | 113.02532 | -1.655408 | 0.1834848 |
| Vermis.X | 0.1426317 | -0.0428193 | 0.3280826 | -7.948177 | 4.952145 | -1.604997 | 0.1903873 |
| Left.VI | 0.1303831 | -0.0513811 | 0.3121473 | -162.09435 | 109.841843 | -1.4757067 | 0.2311349 |
| Vermis.IX | 0.1089864 | -0.0820556 | 0.3000284 | -16.278026 | 13.455817 | -1.2097389 | 0.3525349 |
| Right.I.III | 0.0930177 | -0.0956821 | 0.2817175 | -17.664182 | 17.21659 | -1.0259977 | 0.4496234 |
| Right.VI | 0.0831958 | -0.1005165 | 0.2669081 | -103.01629 | 110.640805 | -0.9310876 | 0.4928105 |
| Vermis.VII | 0.0755074 | -0.1139255 | 0.2649404 | -12.618419 | 15.084266 | -0.8365285 | 0.5373713 |
| Left.V | 0.0715461 | -0.1174341 | 0.2605264 | -32.833845 | 41.665883 | -0.7880271 | 0.5483383 |
| Right.IV | -0.0383012 | -0.2284225 | 0.1518202 | 20.490579 | 48.52677 | 0.4222531 | 0.803883 |
| Left.I.III | 0.0361417 | -0.1528087 | 0.2250921 | -6.424403 | 16.050859 | -0.4002529 | 0.803883 |
| Left.IV | 0.0254305 | -0.1658294 | 0.2166904 | -13.053648 | 46.567167 | -0.2803187 | 0.8727953 |
| Left.X | 0.016973 | -0.1701001 | 0.204046 | -1.395246 | 7.356934 | -0.1896505 | 0.8957806 |
| Right.X | 0.015476 | -0.1731221 | 0.2040741 | -1.251395 | 7.292894 | -0.171591 | 0.8957806 |
| Vermis.VI | 0.0112027 | -0.1749363 | 0.1973417 | -2.544731 | 20.452977 | -0.1244186 | 0.9010054 |

**Table S8.** Effect sizes, betas, standard error and FDR-adjusted P values for cerebellum volume differences between ETLE and healthy controls, controlling for age, age^2^, sex, ICV and site

| Cerebellar region | Cohen's *d* | Lower CI | Upper CI | Beta | Std error | Statistic (*t*) | FDR-adjusted *P* value |
| --- | --- | --- | --- | --- | --- | --- | --- |
| **ETLE** |  |  |  |  |  |  |  |
| Corpus.Medullare | 0.8597582 | 0.6760798 | 1.0434367 | -1709.2888 | 171.128599 | -9.9883293 | **0** |
| Left.VIIB | 0.5174101 | 0.3389493 | 0.6958709 | -532.90335 | 87.601216 | -6.0832871 | **0** |
| Left.VIIIA | 0.4571571 | 0.2789717 | 0.6353426 | -477.95434 | 88.79592 | -5.382616 | **0.0000008** |
| Left.IX | 0.4137427 | 0.2341462 | 0.5933393 | -263.6563 | 53.585886 | -4.9202565 | **0.000007** |
| Right.VIIIB | 0.4090354 | 0.2277486 | 0.5903223 | -251.71838 | 52.795946 | -4.7677596 | **0.0000117** |
| Right.IX | 0.3666018 | 0.191653 | 0.5415507 | -234.46707 | 53.152578 | -4.4112078 | **0.0000481** |
| Right.Crus.II | 0.3554498 | 0.178934 | 0.5319656 | -456.24817 | 102.739549 | -4.4408231 | **0.0000481** |
| Right.VIIB | 0.3314726 | 0.1582739 | 0.5046712 | -396.87758 | 99.689244 | -3.9811474 | **0.0002573** |
| Left.VIIIB | 0.3271309 | 0.1499405 | 0.5043214 | -207.47385 | 53.701361 | -3.8634747 | **0.0003686** |
| Vermis.IX | 0.3042792 | 0.1251935 | 0.4833649 | -45.808125 | 12.671357 | -3.6150924 | **0.0008794** |
| Left.Crus.I | 0.2897334 | 0.1202323 | 0.4592344 | -517.19846 | 146.629847 | -3.5272386 | **0.0011242** |
| Left.Crus.II | 0.2694329 | 0.0932039 | 0.4456618 | -345.40703 | 107.27912 | -3.2197041 | **0.0030859** |
| Right.V | 0.2476886 | 0.0707597 | 0.4246175 | -118.41369 | 40.53434 | -2.9213177 | **0.0076504** |
| Right.Crus.I | 0.2317017 | 0.0585261 | 0.4048774 | -402.94288 | 144.704512 | -2.784591 | **0.0102476** |
| Right.VIIIA | 0.2369443 | 0.0584336 | 0.4154549 | -223.85084 | 80.460789 | -2.7821109 | **0.0102476** |
| Right.X | 0.2105623 | 0.0352879 | 0.3858366 | -17.505216 | 6.992356 | -2.5034791 | **0.0217686** |
| Vermis.X | 0.1949543 | 0.0163424 | 0.3735661 | -10.968091 | 4.704834 | -2.3312384 | **0.0328381** |
| Left.IV | -0.1933479 | -0.3707139 | -0.0159819 | 103.762905 | 45.611519 | 2.2749277 | **0.0359076** |
| Right.I.III | 0.1693399 | -0.0056824 | 0.3443622 | -32.718714 | 16.241325 | -2.0145347 | 0.065131 |
| Left.VI | 0.1555852 | -0.0159094 | 0.3270798 | -197.99168 | 105.189 | -1.882247 | 0.0841799 |
| Vermis.VIII | 0.1132292 | -0.0607713 | 0.2872297 | -32.356554 | 23.998942 | -1.3482492 | 0.2371287 |
| Left.X | 0.1055056 | -0.0679253 | 0.2789365 | -8.645006 | 6.838684 | -1.2641329 | 0.2627866 |
| Left.V | 0.0877487 | -0.0901284 | 0.2656257 | -40.982795 | 39.797558 | -1.0297816 | 0.3692533 |
| Left.I.III | 0.0748509 | -0.0986369 | 0.2483386 | -13.298477 | 14.811597 | -0.8978422 | 0.4310616 |
| Vermis.VI | -0.0677524 | -0.2419371 | 0.1064322 | 15.309701 | 18.934519 | 0.8085603 | 0.469222 |
| Right.VI | -0.0568689 | -0.2273939 | 0.113656 | 72.170386 | 105.502127 | 0.6840657 | 0.5320936 |
| Vermis.VII | 0.0493354 | -0.1259106 | 0.2245814 | -8.323576 | 14.091361 | -0.5906865 | 0.5733918 |
| Right.IV | 0.048153 | -0.1315314 | 0.2278375 | -26.486793 | 47.028036 | -0.5632128 | 0.5733918 |

**Table S9.** Partial η2 effect sizes, beta coefficients, standard error and FDR-adjusted P values for the linear regression of duration of epilepsy against cerebellar volume controlling for age, age2, sex, ICV and site.

| Cerebellar region | partial η^2^ | Lower CI | Upper CI | Beta | Std error | Statistic (*t*) | FDR-adjusted *P* value |
| --- | --- | --- | --- | --- | --- | --- | --- |
| **All epilepsies** |  |  |  |  |  |  |  |
| Total volume | 0.0315631 | 0.0138153 | 0.055591 | -226.7 | 39.49 | -5.741 | **<.0001** |
| Right.VIIIA | 0.0386891 | 0.0128541 | 0.0760663 | -12.90876 | 2.8087212 | -4.5959563 | **0.000121** |
| Right.VIIB | 0.0197437 | 0.0062606 | 0.0401028 | -16.086522 | 3.597154 | -4.4720138 | **0.000121** |
| Left.Crus.II | 0.0194791 | 0.0059392 | 0.040151 | -17.883991 | 4.0979803 | -4.3640987 | **0.0001321** |
| Left.VIIB | 0.0182966 | 0.0054054 | 0.0381844 | -14.421181 | 3.3671011 | -4.2829665 | **0.0001417** |
| Right.VIIIB | 0.0174248 | 0.005025 | 0.0367061 | -8.6848483 | 2.0562328 | -4.2236698 | **0.0001468** |
| Left.VI | 0.0165933 | 0.0044286 | 0.0359231 | -15.939321 | 3.9382174 | -4.0473441 | **0.000261** |
| Right.V | 0.0142428 | 0.0034202 | 0.0319814 | -5.260563 | 1.3703158 | -3.838942 | **0.0005078** |
| Left.V | 0.014059 | 0.0033293 | 0.031712 | -5.3733762 | 1.4089534 | -3.8137358 | **0.0005078** |
| Right.Crus.I | 0.0138153 | 0.0030903 | 0.0317156 | -21.545181 | 5.7986233 | -3.7155682 | **0.0006662** |
| Left.Crus.I | 0.012876 | 0.0026315 | 0.0303762 | -20.706318 | 5.7916984 | -3.5751721 | **0.0009387** |
| Left.VIIIA | 0.0125938 | 0.0025711 | 0.0297292 | -12.13193 | 3.3948033 | -3.5736769 | **0.0009387** |
| Left.VIIIB | 0.0117205 | 0.0021962 | 0.0283566 | -6.8084084 | 1.9705503 | -3.4550797 | **0.0013372** |
| Right.X | 0.0101713 | 0.0015691 | 0.0258953 | -0.7951659 | 0.2463054 | -3.228374 | **0.002607** |
| Right.Crus.II | 0.0107127 | 0.0016467 | 0.0272755 | -13.553534 | 4.2030055 | -3.2247243 | **0.002607** |
| Right.VI | 0.0100268 | 0.0014928 | 0.0257458 | -12.312375 | 3.8569655 | -3.1922439 | **0.0027169** |
| Vermis.VII | 0.0091135 | 0.0011614 | 0.0242506 | -1.6074644 | 0.5270395 | -3.0499886 | **0.0041092** |
| Right.IV | 0.0081675 | 0.0008475 | 0.022665 | -4.939581 | 1.7062201 | -2.8950433 | **0.0063775** |
| Left.X | 0.0090238 | 0.0005157 | 0.0273347 | -0.6493216 | 0.2502501 | -2.5946904 | **0.0150193** |
| Left.I.III | 0.0063268 | 0.0002806 | 0.0197994 | -1.2992463 | 0.5171612 | -2.5122659 | **0.0179108** |
| Corpus.Medullare | 0.0059048 | 0.0002052 | 0.0188874 | -15.770055 | 6.4315343 | -2.4519895 | **0.019544** |
| Left.IX | 0.0062586 | 0.0002105 | 0.0200533 | -4.9634117 | 2.0298595 | -2.4451996 | **0.019544** |
| Right.I.III | 0.0056527 | 0.0001469 | 0.0184482 | -1.3233447 | 0.5518788 | -2.3978901 | **0.0212167** |
| Vermis.VIII | 0.0054619 | 0.0001084 | 0.0180829 | -2.1520631 | 0.9116859 | -2.3605314 | **0.0224452** |
| Vermis.IX | 0.003934 | 0 | 0.0152781 | -0.9402057 | 0.4707327 | -1.997324 | 0.0537342 |
| Right.IX | 0.0037601 | 0 | 0.0151608 | -3.7842166 | 1.9672124 | -1.9236441 | 0.0612511 |
| Left.IV | 0.0024761 | 0 | 0.0122251 | -2.6227091 | 1.6491394 | -1.5903501 | 0.1206866 |
| Vermis.VI | 0.0008512 | 0 | 0.0086889 | -0.6164964 | 0.6948728 | -0.8872075 | 0.3890943 |
| Vermis.X | 0.0003868 | 0 | 0.0064859 | -0.109424 | 0.1745248 | -0.6269824 | 0.5308116 |
| **TLE-HS-L** |  |  |  |  |  |  |  |
| Total volume | 0.0088775 | 0 | 0.052985 | -137.8 | 105.3 | -1.309 | 0.3733333 |
| **TLE-HS-R** |  |  |  |  |  |  |  |
| Total volume | 0.0644833 | 0.0077235 | 0.1596294 | -311.1 | 103.3 | -3.012 | **0.0095455** |
| **TLE-NL-L** |  |  |  |  |  |  |  |
| Total volume | 0.02806 | 0 | 0.1234849 | -198.5 | 121.8 | -1.629 | 0.2340625 |
| **TLE-NL-R** |  |  |  |  |  |  |  |
| Total volume | 0.0743044 | 0.0011834 | 0.2129315 | -304.2 | 127.7 | -2.382 | 0.0538462 |
| **GGE** |  |  |  |  |  |  |  |
| Total volume | 0.007365 | 0 | 0.0558724 | -125.1 | 116.3 | -1.075 | 0.497 |
| **ETLE** |  |  |  |  |  |  |  |
| Total volume | 0.0543002 | 0.0062654 | 0.1373488 | -303.4 | 101.8 | -2.981 | **0.0095455** |

**Table S10.** Partial η2 effect sizes, beta coefficients, standard error and FDR-adjusted P values for the linear regression of age of epilepsy onset against cerebellar volume controlling for age, age2, sex, ICV and site

| Cerebellar region | partial η^2^ | Lower CI | Upper CI | Beta | Std error | Statistic (*t*) | FDR-adjusted *P* value |
| --- | --- | --- | --- | --- | --- | --- | --- |
| **All epilepsies** |  |  |  |  |  |  |  |
| Total volume | 0.0244551 | 0.0090364 | 0.0466015 | 196 | 39.42 | 4.972 | **<.0001** |
| Right.VIIB | 0.0163431 | 0.004375 | 0.0353544 | 14.4992811 | 3.5765947 | 4.0539346 | **0.0015209** |
| Right.Crus.I | 0.0137782 | 0.0030085 | 0.0318586 | 21.5312531 | 5.8568052 | 3.6762796 | **0.0024071** |
| Left.VIIB | 0.013868 | 0.0030125 | 0.0321115 | 12.3491436 | 3.366697 | 3.6680294 | **0.0024071** |
| Left.VI | 0.0133284 | 0.0026101 | 0.0318004 | 13.9173452 | 3.9601353 | 3.514361 | **0.0029301** |
| Left.Crus.I | 0.0122056 | 0.0022673 | 0.0295804 | 20.0072064 | 5.8097538 | 3.4437271 | **0.0029301** |
| Left.Crus.II | 0.0116755 | 0.0021441 | 0.0284008 | 14.2033075 | 4.1421017 | 3.4290098 | **0.0029301** |
| Right.VIIIA | 0.0122197 | 0.0021281 | 0.0301139 | 9.8214123 | 2.9200349 | 3.3634572 | **0.0029301** |
| Right.V | 0.0110423 | 0.0019019 | 0.0273317 | 4.6465525 | 1.386896 | 3.350325 | **0.0029301** |
| Left.V | 0.0107032 | 0.0017641 | 0.0267977 | 4.7098221 | 1.4273492 | 3.2996985 | **0.0031168** |
| Right.VIIIB | 0.0105125 | 0.0015997 | 0.026837 | 6.5567846 | 2.0398481 | 3.2143494 | **0.0037813** |
| Left.VIIIA | 0.0014162 | 0.0258455 | 10.6983302 | 3.3987995 | 3.1476792 | 0.0043151 | **0.0099598** |
| Right.VI | 0.0097315 | 0.0013453 | 0.025421 | 12.1697436 | 3.8988126 | 3.1213974 | **0.0043215** |
| Right.X | 0.0092507 | 0.001173 | 0.024636 | 0.7528699 | 0.2471801 | 3.0458355 | **0.0051295** |
| Right.Crus.II | 0.007203 | 0.0004248 | 0.0218167 | 10.9392196 | 4.1983766 | 2.6055832 | **0.0186349** |
| Right.IV | 0.0056137 | 0.0001321 | 0.0184242 | 4.1399692 | 1.737358 | 2.3829109 | **0.0324076** |
| Vermis.VII | 0.0091135 | 0.0011614 | 0.0242506 | -1.6074644 | 0.5270395 | -3.0499886 | **0.0041092** |
| Right.IV | 0.0081675 | 0.0008475 | 0.022665 | -4.939581 | 1.7062201 | -2.8950433 | **0.0063775** |
| Left.VIIIB | 0.0056132 | 0.0000846 | 0.0187798 | 4.5687926 | 1.9590609 | 2.332134 | **0.0348231** |
| Vermis.VII | 0.0051731 | 0.000028 | 0.0177128 | 1.2000389 | 0.5273604 | 2.2755573 | **0.0380212** |
| Corpus.Medullare | 0.0048988 | 0 | 0.0171768 | 14.3793214 | 6.4807368 | 2.2187788 | **0.0415732** |
| Left.X | 0.005544 | 0 | 0.0201669 | 0.5362937 | 0.2518611 | 2.1293232 | **0.0494087** |
| Left.I.III | 0.0044602 | 0 | 0.0168441 | 1.0601447 | 0.5164126 | 2.0529025 | 0.0565015 |
| Vermis.VIII | 0.0032226 | 0 | 0.0139115 | 1.6461056 | 0.914016 | 1.8009593 | 0.091826 |
| Right.I.III | 0.0032549 | 0 | 0.0140562 | 0.9842423 | 0.546772 | 1.8000965 | 0.091826 |
| Left.IV | 0.0025194 | 0 | 0.0124088 | 2.6489874 | 1.6622041 | 1.5936595 | 0.135528 |
| Left.IX | 0.0018124 | 0 | 0.0113748 | 2.5920419 | 2.0061214 | 1.2920663 | 0.2294353 |
| Right.IX | 0.001258 | 0 | 0.0095918 | 2.1412015 | 1.9367741 | 1.1055504 | 0.3014986 |
| Vermis.IX | 0.0006381 | 0 | 0.0075757 | 0.3762742 | 0.4726042 | 0.7961719 | 0.4589011 |
| Vermis.VI | 0.0003948 | 0 | 0.0069122 | 0.4274503 | 0.7022036 | 0.608727 | 0.5629584 |
| Vermis.X | 0.0000032 | 0 | 0.0011967 | 0.0100437 | 0.1769575 | 0.0567578 | 0.9547496 |
| **TLE-HS-L** |  |  |  |  |  |  |  |
| Total volume | 0.0027339 | 0 | 0.0358659 | 72.75 | 99.87 | 0.728 | 0.6538 |
| **TLE-HS-R** |  |  |  |  |  |  |  |
| Total volume | 0.0726441 | 0.0081317 | 0.1796213 | 304.8 | 102.8 | 2.964 | **0.0155556** |
| **TLE-NL-L** |  |  |  |  |  |  |  |
| Total volume | 0.0006938 | 0 | 0.0473568 | 31.84 | 132.6 | 0.24 | 0.8870313 |
| **TLE-NL-R** |  |  |  |  |  |  |  |
| Total volume | 0.0263673 | 0 | 0.131895 | 171.5 | 119.3 | 1.438 | 0.3170588 |
| **GGE** |  |  |  |  |  |  |  |
| Total volume | 0.005655 | 0 | 0.0520448 | 110 | 118.2 | 0.93 | 0.59 |
| **ETLE** |  |  |  |  |  |  |  |
| Total volume | 0.0504851 | 0.0047312 | 0.1327793 | 303 | 106.6 | 2.843 | **0.0175** |

**Epilepsy syndrome comparison analyses**

Additional analyses were conducted to compare epilepsy syndromes. Consistent with the main analyses, we fit linear mixed models to investigate differences in total and regional cerebellar volume with age, sex and ICV modelled as fixed factors and site as a random factor. Given the significant discrepancies in duration of illness between the lesional (TLE-HS) and non-lesional (TLE-NL) syndromes as well as differences in age between some syndromes (see Table 2), duration of illness was modelled as a fixed effect in all models. The following comparisons were made:

TLE-HS-L and TLE-NL-L

TLE-HS-R and TLE-NL-R

GGE and TLE-HS-L

GGE and TLE-HS-R

Extra temporal focal epilepsy and GGE

Extra temporal focal epilepsy and TLE-HS-L

Extra temporal focal epilepsy and TLE-HS-R

Extra temporal focal epilepsy and TLE-NL-L

Extra temporal focal epilepsy and TLE-NL-R

Results were FDR (*P*<0.05) corrected for multiple comparisons.

**Secondary analyses**

We assessed cerebro-cerebellar relationships to assess whether the magnitude of cerebellar tissue changes observed in people with epilepsy, mirrored that seen in the cerebral cortex. For this analysis, we used Freesurfer-derived cortical thickness measures of the left and right hemisphere. A ‘total’ cortical thickness measure was obtained by calculating the average of the left and right hemisphere cortical thickness measures, for each individual. We fit the following linear mixed model:

**Total vol ~ Cerebral cortical thickness * Dx + Sex + Age + Age(^2) + ICV + (1| Site)**

where total volume is predicted by the interaction between cerebral cortical thickness and diagnosis.

Using this model, results showed a significant interaction between cortical thickness and diagnosis on predicting cerebellar total volume (p=0.01; controlling for all other covariates). Marginal effects plots showed that a reduction in mean cortical thickness was associated with a more pronounced decrease in cerebellar volume for those with epilepsy (Figure S2).

Additionally we explored the interaction of subdiagnosis group with cortical thickness in a model:

**Total vol ~ Cerebral cortical thickness * SDx.group +  Sex + Age + Age(^2) + ICV + (1| Site)**

where SDx.group is TLE-MTS, TLE non-lesional, GGE and Extratemporal focal epilepsy. Left and right focal groups were combined within each subtype.

While all epilepsies other than TLE non-lesional showed a similar trend, only TLS-MTS showed a statistically significant interaction between cortical thickness and diagnosis (p=0.003), suggesting that the reduction in cerebellum volume relative to cortical thickness is more pronounced in lesional epilepsies.

**Table S11** Clinical characteristics of people with epilepsy treated and never treated, with Phenytoin.

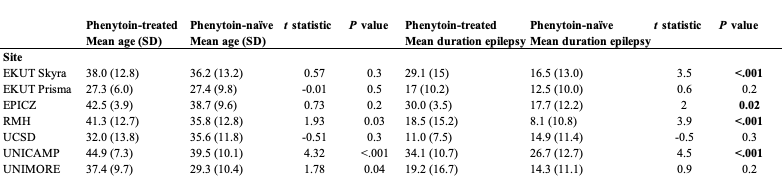

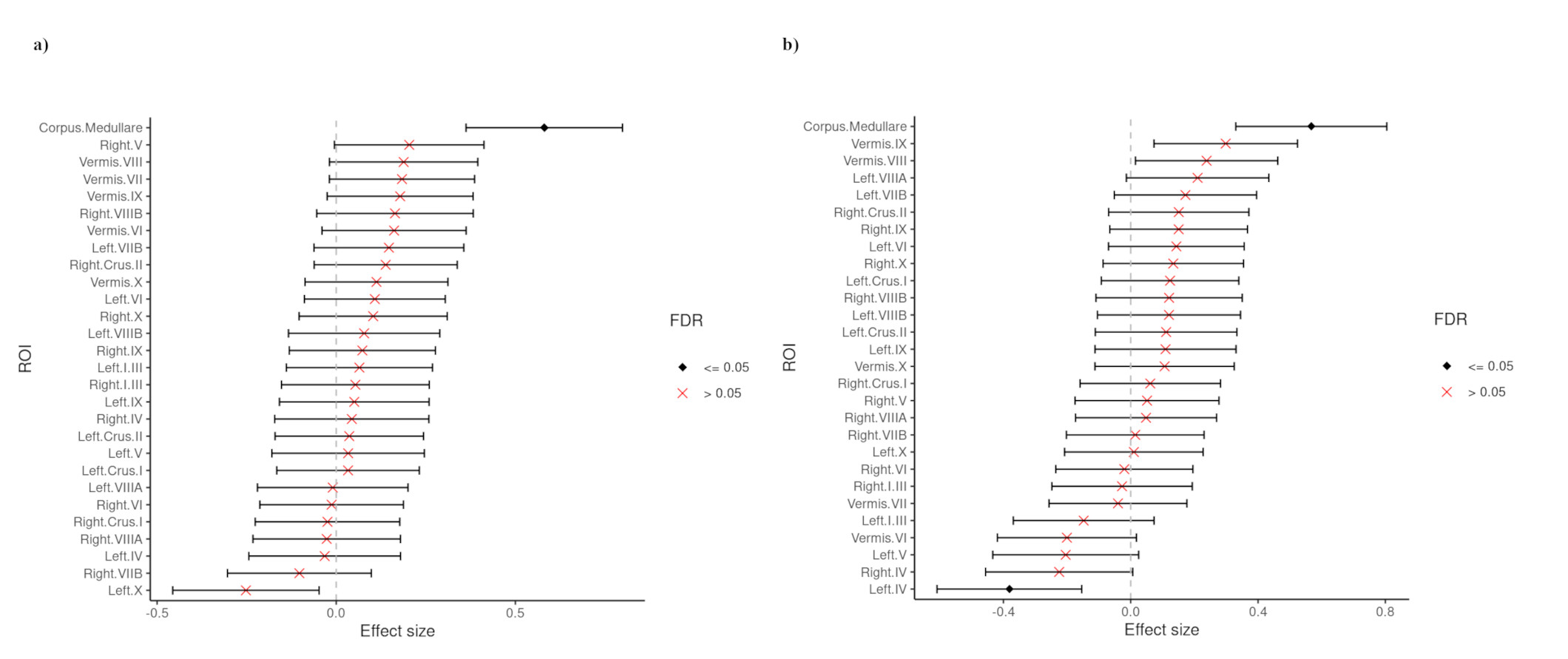

**Figure S1.** Forest plots (Cohen’s *d* +/- 95% confidence interval) showing the between-group differences for a) TLE-NL-L and b) TLE-NL-R epilepsy vs. healthy controls (HC).

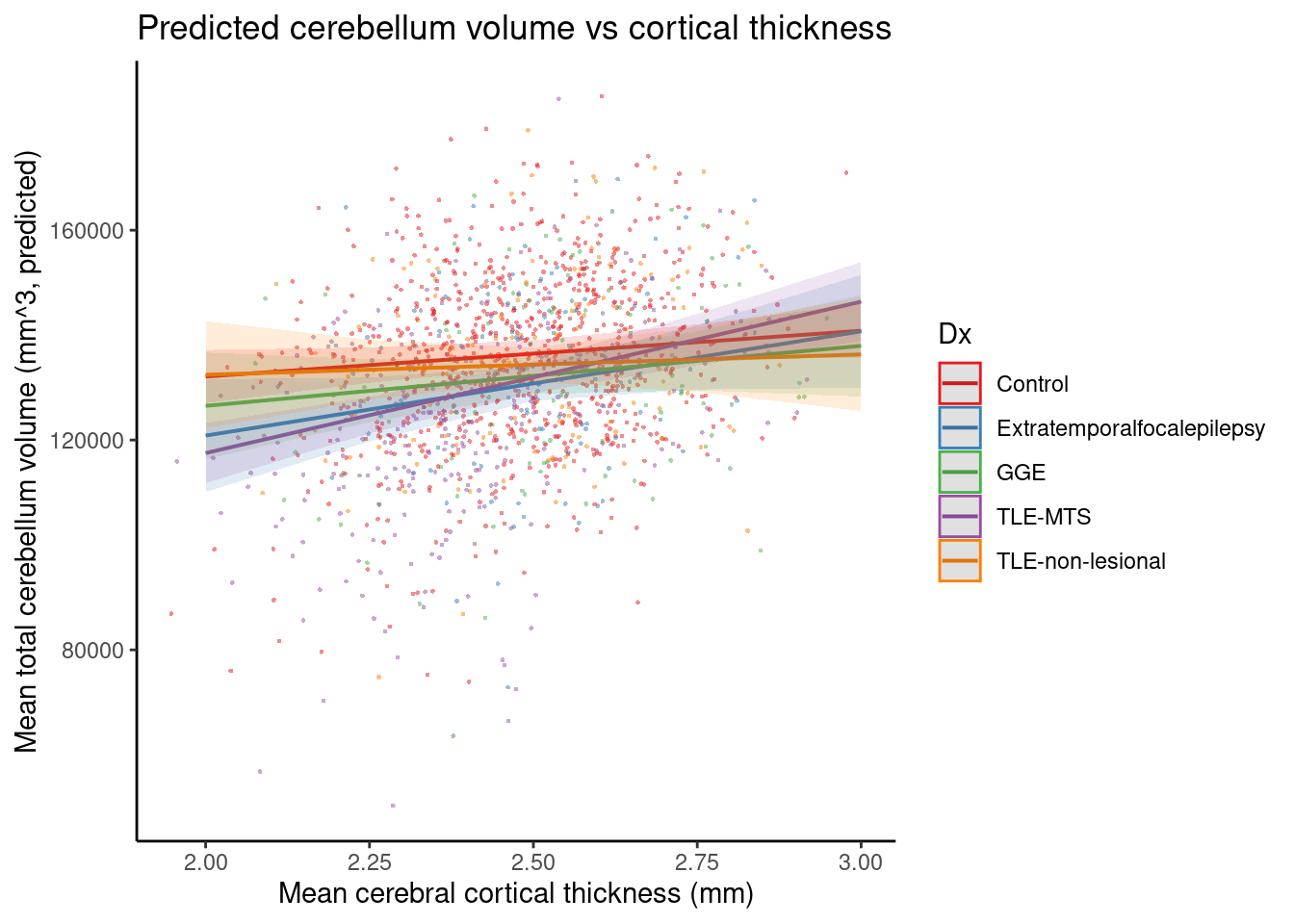

**Controls**

**ETLE**

**GGE**

**TLE-HS**

**TLE-NL**

**Figure S2.** Marginal effects plot (controlling for all covariates) showing the relationship between total cerebellar volume and cortical thickness, in controls (red) and each of the epilepsy syndromes. For individuals at the lower extremes of cortical thickness with a mean of 2.25 mm, this amounts to a difference in mean total cerebellar volume of 7832 [CI: 7602 - 8062] mm^3^ between cases and controls.

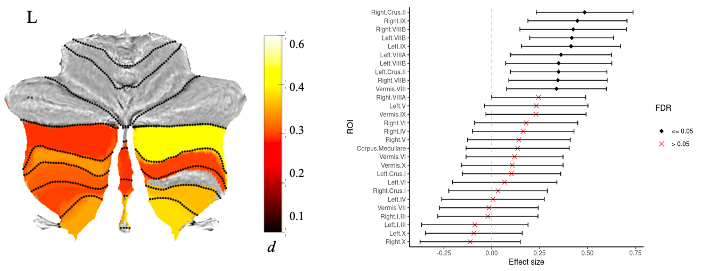

**I-IV**

**V**

**VI**

**Crus I**

**Crus II**

**VIIB**

**VIIIA**

**IX**

**X**

**VIIIB**

**Figure S3.** Atlas-based effect size (Cohen’s *d*) maps, MNI-based coronal slices (top: y= -72; bottom: y= -54) and forest plots (Cohen’s *d* +/- 95% confidence interval) of the significant association between Phenytoin use and cerebellar regional volume, in epilepsy. Positive effect sizes reflect people with epilepsy treated with Phenytoin < people with epilepsy never treated with Phenytoin. Regions significant at *P* < 0.05 FDR corrected are depicted.

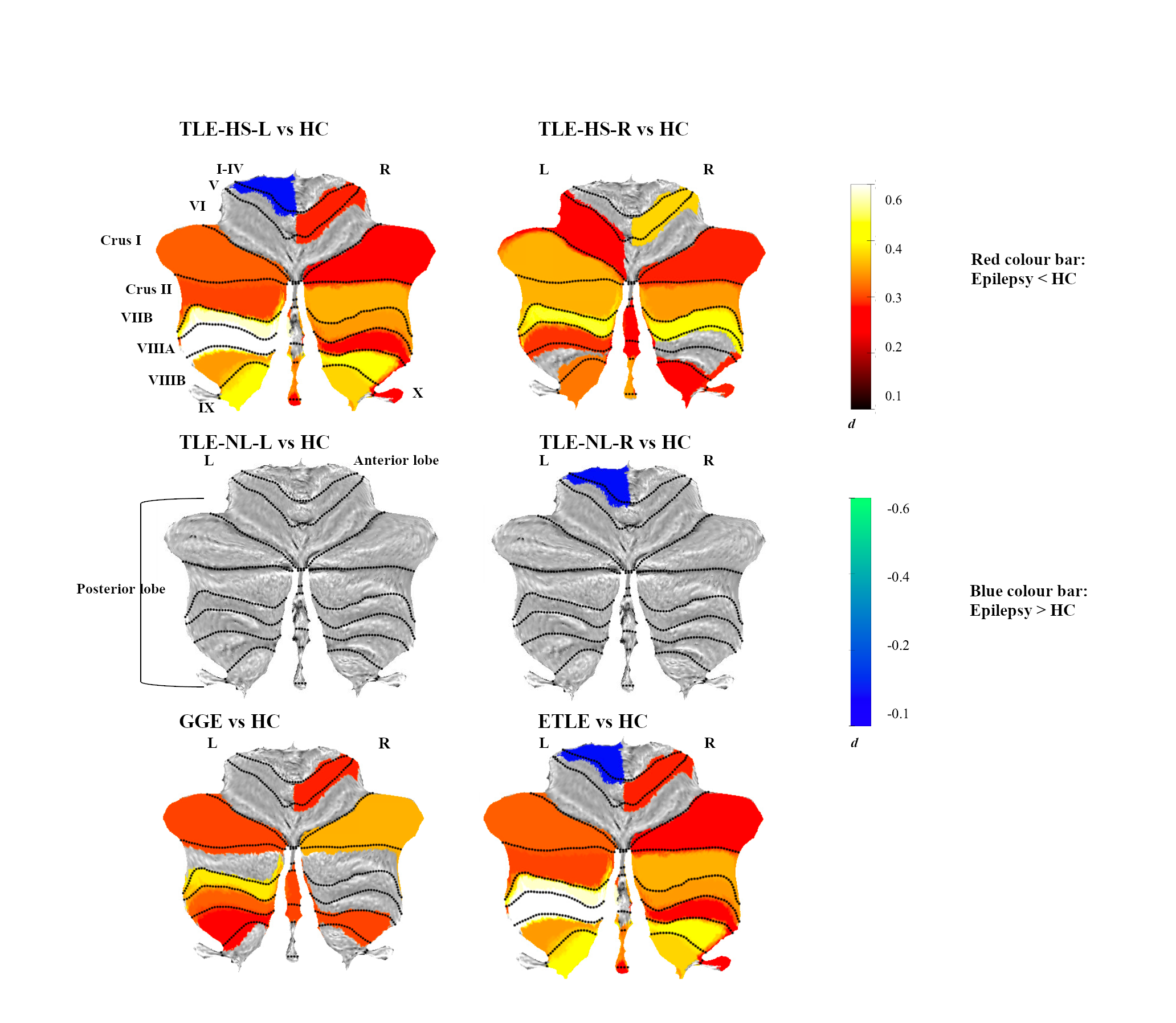

**Figure S4.** Summary atlas-based effect size (Cohen’s *d*) map showing the main effect sizes for each of the epilepsy syndrome (versus healthy controls; HC) comparisons. Top row L to R: TLE-HSL vs HC, TLE-HS-R vs HC. Middle row L to R: TLE-NL-L vs HC, TLE-NL-R vs HC. Bottom row L to R: GGE vs HC, ETLE vs HC. Positive effect sizes (represented by red-yellow colour) reflect cerebellar regions that were smaller in epilepsy compared to HC. Negative effect sizes (represented by blue-green colour) reflect cerebellar regions that were larger in epilepsy compared to HC. All regions shown were significant at *P <* 0.05 FDR corrected.
